# PAC/SP3 on-bead carboxyl derivatization allows combined C- and N-terminomics

**DOI:** 10.1101/2025.09.28.679096

**Authors:** Kristian I. Karlic, Alexander R. Ziegler, Laura E. Edgington-Mitchell, Nichollas E. Scott

**Author notes:** To whom correspondence and requests for materials should be addressed to N.E.S and L.E.E.M. These authors contributed equally to this publication (Kristian I. Karlic, Alexander R. Ziegler).

## Abstract

On-bead single-pot solid-phase enhanced sample preparation, SP3, also known as Protein Aggregation Capture (PAC), is a robust, high-throughput, and widely utilized approach for proteomic sample preparation. Recent studies have highlighted PAC/SP3 as an ideal platform for chemoproteomics, allowing chemical labelling by minimizing sample loss and improving recovery of derivatized peptides. In this work, we establish an on-bead PAC/SP3 protein-level amine and carboxyl derivatization approach to facilitate C-terminal focused proteomics. We demonstrate that on-bead protein derivatization of carboxyl groups can be achieved using ethanolamine, (2-aminoethyl)trimethylammonium (AETMA), and (carboxymethyl)trimethylammonium (Girard’s reagent T, GT) via EDC/HOBt coupling, enabling the labelling of protein C-termini. Using a prokaryotic model system, *Acinetobacter baumannii*, we demonstrate that AETMA and ethanolamine labelling each enables the identification of unique protein C-terminal peptides, with AETMA improving the identification of C-terminal peptides lacking basic residues. Finally, we apply this approach to interrogate both N- and C-termini in response to etoposide-induced apoptosis within Jurkat cells, demonstrating that combined N- and C-terminomics is achievable using on-bead derivatization, yet provides modest coverage of the C-terminome in its current form. Overall, this work establishes bead-based carboxyl group derivatization as a viable platform to enable future C-terminomics method development.

## Introduction

LC-MS-based proteomics is an indispensable tool for the analysis of protein samples and the identification of protein modifications. Over the last 20 years, significant progress has been made in the refinement of sample preparation techniques (1, 2), data collection approaches (3, 4), and bioinformatic tools (5, 6). While current proteomic approaches allow in-depth characterization of proteomes, not all peptides are equally amenable to analysis using standard positive mode LC-MS. Dramatic differences in ionization efficiency and peptide fragmentation properties directly impact the diversity of peptides routinely identified across proteomic experiments (7, 8). While the use of alternative proteases and/or alternative fragmentation approaches has been demonstrated to expand the observable proteome (9–12), many peptides, such as those derived from the C-termini of proteins, remain recalcitrant to standard LC-MS workflows. A key challenge associated with the analysis of C-terminal peptides, referred to as C-terminomics, is the lack of basicity within these peptides if tryptic digestion has been used during sample preparation. The generation of these so called “base-less” peptides has been shown to result in precursors that exhibit low charge density, which can result in their exclusion from selection if only observed as singly-charged species (13). Additionally, peptides lacking basic residues may result in less informative spectra, as their fragmentation patterns can lack complementary ion series typically observed within standard tryptic peptides in positive ion mode (8, 14). To combat these limitations, robust and accessible approaches to improve both the detection and identification of the C-terminome are required, with chemical derivatization offering an attractive solution.

Chemical derivatization of peptides is widely used to expand the capabilities of LC-MS-based proteomic analysis. Derivatization approaches can be used to improve peptide ionization efficiency (15), augment peptide fragmentation properties (16–20), and permit sample multiplexing (21–25). In the context of degradomics, the labelling of protein termini prior to digestion is essential to facilitate N-(26–28), and C-(29–32) terminomics studies, as the introduction of labels provides chemical proof of the position of amino acids within the intact polypeptide before digestion. While robust and high-efficiency amine labelling approaches are available, such as reductive dimethylation (21), approaches for C-terminal protein labelling are less refined. To date, C-terminal proteome studies have utilized carbodiimide chemistry, wherein 1-ethyl-3-(3-dimethylaminopropyl) carbodiimide (EDC) and the activators N-hydroxysuccinimide (NHS) or 1-hydroxy-7-azabenzotriazole (HOAt) are used to modify carboxylic acid with agents such as 2-aminoethyl trimethylammonium (AETMA), carboxymethyl trimethylammonium (Girard’s reagent T, GT), or ethanolamine (18, 29, 31–33). The modification of carboxylic acids with these labels has been shown to both increase the charge states of modified peptides as well as improve the identification of C-terminal peptides (16, 18, 32). However, a key issue noted in prior C-terminomic studies has been ensuring protein solubility, with high concentrations of chemically inert chaotropic agents such as guanidinium chloride (29, 30) previously needed to be employed. These reagents can be incompatible with downstream proteomic approaches, requiring additional sample handling to reduce or remove these agents, limiting the versatility of C-terminal labelling.

Protein Aggregation Capture (PAC) and Single-Pot Solid-Phase-Enhanced Sample Preparation (SP3) are an increasingly utilised family of proteomics preparation approaches (34, 35). These sample preparation approaches utilize on-bead protein precipitation with organic solvents to allow the removal of contaminants, providing a highly effective and automatable method for sample preparation (34, 35). While initial adoption of PAC/SP3 was driven by the versatility and simplicity of these approaches, several teams have highlighted the unique capacity for PAC/SP3 to streamline cumbersome and loss-prone sample clean-up steps. This is best illustrated by the work of the Backus lab, who demonstrated the capacity for PAC/SP3 to significantly improve chemoproteomic studies of cysteines by eliminating undesirable reagents detrimental to downstream analysis (36–38). Within the field of N-terminomics, PAC/SP3 has been shown to be an ideal platform to allow on-bead N-terminal-based derivatization with both reductive dimethylation (39) and NHS-based isobaric labelling (40) reported with these platforms to date, having also shown improvements in throughput via automation (41). We have previously utilized on-bead N-terminal derivatization to allow N-terminome analysis of the cysteine protease legumain across multiple tissues (28). By combining N-terminal derivatization with peptide fractionation or data-independent acquisition (42, 43) we have demonstrated that both N-terminomic and proteomic analyses can be achieved within a single experiment. However, to allow true global degradomics of a given sample, the integration of both N- and C-terminomics would be highly advantageous.

Within this study, we demonstrate the proof-of-principle capacity of on-bead carboxyl derivatization to allow C-terminomics analysis. Using simple (avidin) and complex (*Acinetobacter baumannii* and apoptotic human T-lymphocytes) samples, we assess the performance of on-bead carboxyl group derivatization with ethanolamine, AETMA, and Girard’s T Reagent for the identification of protein C-termini. We demonstrate that sequential labelling enables the successful installation of C-terminal labels within proteome samples, and that different C-terminal labelling reagents allow the identification of discrete subsets of C-terminal peptides without the need for enrichment of C-terminal peptides. Combined, this work demonstrates the capacity of bead-based derivatization of carboxyl groups to allow total proteomic analysis and combined N-/C-terminomics within a single sample.

## Methods

### Preparation of bacterial cell lysates

*A. baumannii* D1279779 lysates were prepared as previously described with minor modifications (44, 45). Briefly, *A. baumannii* D1279779 (four biological replicates) was grown to stationary phase overnight in Luria broth, then washed 3 times in chilled PBS (137 mM NaCl, 2.7 mM KCl, 10 mM Na_2_HPO_4_, and 1.8 mM KH_2_PO_4_) before being boiled in SDS lysis buffer (4% SDS, 100 mM HEPES pH 8.0) at 95°C with shaking for 10 min. Protein concentrations were then determined by bicinchoninic acid assays (Thermo Fisher Scientific).

### Preparation of Avidin

Avidin (Sigma Aldrich - A9275) was resuspended in SDS lysis buffer, boiled and 50 µg aliquots were prepared. All avidin experiments were undertaken in technical triplicate on independent aliquots of avidin.

### Cell culture and induction of apoptosis in Jurkat cells

Jurkat cells (human T lymphocytes, ATCC TIB-152) were cultured in RPMI medium supplemented with glutamine (Gibco, 11875093), 10% foetal bovine serum (FBS, CellSera) and 1% antibiotics (100 U/mL penicillin-streptomycin, Thermo Fisher Scientific) at 37°C with 5% CO_2_. Cells were passaged 1:10. To induce apoptosis, 2×10^6^ Jurkat cells were seeded in a 6-well plate and grown overnight. Cell media was replaced, and 50 µg/mL etoposide (Sigma #E1383) added to cells for 8 hours at 37°C with 5% CO_2_ with an equal volume of DMSO added as a vehicle control. Live cell labelling of caspases was achieved by adding the activity-based probe LE22 (46) to cells 30 min prior to harvesting. Cells were harvested by centrifugation (300 *x g* for 3 min) and washed in PBS. For gel-based assays, cells were lysed in PBS containing 0.1% Triton X-100 (PBSTX, Sigma) supplemented with Roche cOmplete EDTA-free protease inhibitor (Roche, 11836170001) on ice. Cell debris was removed by centrifuging at 21,000 *x g* for 7 minutes at 4°C and protein amount was normalised by bicinchoninic acid (BCA) assay per manufacturer’s instructions (Thermo). A total of 50 µg protein per sample was diluted in 20 µL PBSTX and 5x sample buffer (50% glycerol, 250 mM Tris pH 6.8, 10% SDS, 0.04% bromophenol blue, 6.25% beta-mercaptoethanol) was added. Samples were then boiled for 5 min and loaded onto a freshly prepared 15% acrylamide gels for SDS-PAGE. Gels were scanned for Cy5 fluorescence using a Typhoon 5 flatbed laser scanner (GE Healthcare) to visualise caspase labelling. Proteins were then transferred to a nitrocellulose membrane using the Trans-Blot Turbo Transfer system (Biorad) and stained with Ponceau S. All antibodies were diluted in 1:1 intercept blocking buffer (Li-Cor) and PBS containing 0.05% Tween-20 (PBST; Sigma). Membranes were incubated with rabbit anti-caspase 3 (1:1,000; Abcam ab2302) overnight at 4°C followed by detected with goat anti-rabbit IgG HRP (1:10,000; Abcam ab6721), which was incubated for 60 min at room temperature. Chemiluminescence was detected by adding Clarity ECL Substrate (Biorad) and scanning on a ChemiDoc (BioRad).

### Preparation of cell lysates

Cells were seeded and treated with etoposide to induce apoptosis as indicated above, without the addition of LE22. Following washing of cell pellets in PBS, cells were lysed in SDS lysis buffer supplemented with Roche cOmplete EDTA-free protease inhibitor. Lysis was facilitated by two rounds of probe sonication for 3 sec (30% amplitude), followed by boiling for 10 minutes. Lysates were cleared to remove cell debris by centrifugation (21,000 *x g* for 7 min, 4°C) and normalised by BCA assay. Samples were diluted in SDS lysis buffer (50 µg in 100 µL) and stored at - 20°C until proteomic analysis.

### On-bead derivatization

Amine derivatization was undertaken utilizing the approach of Weng et al. (31471496). Briefly, 50 µg of samples (Avidin, bacterial, or cell lysate) in SDS lysis buffer were reduced with 10 mM TCEP and alkylated with 40 mM CAA for 30 min at 45°C with 1500 rpm in the dark. Following reduction/alkylation, equal volumes of carboxylate-functionalized beads (Sera-Mag Speed Beads, GE Life Sciences, cat. no. 45152105050350 and 65152105050350) were washed twice with water prior to addition at a 10:1 bead:protein ratio in a final concentration of 80% v/v ethanol. Samples were incubated at room temperature for 10 min at 1,000 rpm to initiate binding and were washed twice with 80% v/v ethanol. Protein amine groups were then protected on-bead via reductive dimethylation (39) with 30 mM formaldehyde and 30 mM sodium cyanoborohydride in 200 mM HEPES pH 7.5 for 1 hr at 37°C, 1,500 rpm, with additional fresh labelling reagents added for another hour as described above to ensure complete labelling. Additional beads were then added at a 5:1 ratio, and protein was bound by the addition of ethanol to a final concentration of 80% v/v ethanol for 10 min at 1,000 rpm to ensure protein binding before being washed twice with 80% v/v ethanol.

Carboxyl derivatization was undertaken based on the protocol of Solis et al. (30). To allow carboxyl derivatization of dimethylated proteins samples were first resuspended in 400 mM MES (pH 5.0) and 160 mM EDC (1-Ethyl-3-[3-dimethylaminopropyl]carbodiimide hydrochloride - Thermo Fischer Scientific – 22980 prepared in 85% DMSO), followed by 160 mM HOBt (1-Hydroxybenzotriazole hydrate - Sigma Aldrich – 54802 prepared in DMSO) added to samples before being incubated at 25°C for 30 min, 1,100 rpm. For carboxyl group labelling AETMA (2-Aminoethyl)trimethylammonium chloride hydrochloride - Sigma Aldrich - 284556), Girard’s T Reagent - Sigma Aldrich - G900) or ethanolamine (Sigma Aldrich – 15014) were prepared in DIPEA (*N*,*N*-Diisopropylethylamine - Sigma Aldrich – 496219) at equivalent amounts and added at a concentration of 240 mM and incubated for 1 hr at 25°C, 1,100 rpm. Fresh EDC/HOBt/carboxyl group labelling reagents were added as above an hour later, then left to incubate overnight at 25°C, 1,100 rpm. To improve labelling efficiency, additional labelling was undertaken by recollecting proteins by adding beads at a 5:1 ratio then adding 80% v/v ethanol before shaking for 10 minutes at 1,100 rpm then repeating carboxyl group labelling as described above. The following day carboxyl derivatised samples were collected on bead by the addition of 80% v/v ethanol for 10 minutes at 1,100 rpm before being washed 3 times with 80% v/v ethanol. Avidin samples were then digested with Pepsin resuspended in 100mM glycine-HCl pH 1.6 (1:50 w/w, Promega) while *A. baumannii* and Jurkat samples were digested with Trypsin in 100 mM HEPES pH 8.0 (1/100 w/w, Promega). All digests were incubated for 16-18 hr at 37°C, 1,000 rpm on a Thermomixer. Following digestion, supernatants were collected using a magnetic rack, acidified to a final concentration of 1% formic acid and desalted on C18 StageTips as previously described (47). StageTips were washed with 10 bed volumes of Buffer B (0.1% formic acid, 80% acetonitrile), then equilibrated with 10 bed volumes of Buffer A* (0.1% trifluoroacetic acid (TFA), 2% acetonitrile) before use. Samples were loaded on to equilibrated columns which were washed with at least 10 bed volumes of Buffer A* before bound peptides were eluted with Buffer B. Eluted peptides were dried by vacuum centrifugation and stored at −20 °C.

### Protein LC-MS analysis

To assess labelling efficiency at the protein level, labelled Avidin was eluted from beads according to the approach of Dagley et al. (48). Briefly, upon completion of labelling, −20 °C formic acid was added to beads and vigorously vortexed for 30 sec, followed by centrifugation at 10,000 *x g* for 1 min. Using a magnetic rack, released proteins was collected and the formic acid neutralised by the addition of 1M Triethylammonium bicarbonate pH 8.5 (Sigma). Protein samples were analysed on a Waters Synapt XS QTof using a Jupiter 300 C5 column (2mm*50mm, Phenomenex). Protein samples were loaded directly onto the C5 column at a flow rate of 0.25 mL/min. Two μg of protein was desalted on column for 2 min with Buffer A (2% acetonitrile, 0.1% formic acid) before being separated by altering the percentage of Buffer B (80% acetonitrile, 0.1% formic acid) from 0% to 100% over 16.5 min. The column was then held at 100% Buffer B for 0.5 min before being equilibrated for 1 min with Buffer A, for a total run time of 20 min. Samples were then deconvoluted with UniDec (v.7.0.0b) (49) (Settings: m/z: 600-2,200; Charge Range: 6-26; Mass Range: 13,500-18,000 Da, Sample Mass: 1 Da, Smooth ChargeStates Distributions, “Some” for Smooth Nearby points and Supress Artifacts, and Peak Dectection Range of 12 with a detection threshold of 0.1).

### Proteomic LC-MS/MS analysis

Proteomic samples were resuspended in Buffer A* (2% acetonitrile, 0.1% TFA) and separated using a two-column chromatography set up composed of a PepMap100 C18 20Lmm × 75 μm trap and a PepMap C18 500Lmm × 75 μm analytical column (Thermo Fisher Scientific). Samples were concentrated onto the trap column at 4-5LμL/min for EThcD samples for 6 min with Buffer A (0.1% formic acid, 2% DMSO) and then infused into an Orbitrap Fusion Lumos Tribrid Mass Spectrometer (Thermo Fisher Scientific) at 300LnL/min via the analytical column using a Dionex Ultimate 3000 UPLC (Thermo Fisher Scientific). Avidin proteomics samples were analysed using identical chromatography conditions using 96-min analytical runs by altering the buffer composition from 2% Buffer B (0.1% formic acid, 77.9% acetonitrile, 2% DMSO) to 28% B over 59 min, then from 28% B to 40% B over 10 min, then from 40% B to 80% B over 7 min. The composition was held at 80% B for 3 min, and then dropped to 2% B over 1 min before being held at 2% B. The Lumos Mass Spectrometer was operated in a data-dependent acquisition mode acquiring a single Orbitrap MS scan (450–2,000Lm/z, maximal injection time of 246 ms, an Automated Gain Control (AGC) set to a maximum of 100% and a resolution of 120k) was collected every 3 s followed by a Orbitrap MS/MS HCD/CID/EThcD scans of precursors (HCD: NCE 15%, stepped 10% two times, maximal injection time of 250Lms, an AGC set to a maximum of 500%L and a resolution of 30k; CID: NCE 30%, maximal injection time of 250 ms, an AGC set to a maximum of 500%L and a resolution of 30k; EThcD: 120–6,000Lm/z, maximal injection time of 250 ms, an Automated Gain Control (AGC) set to a maximum of 500% and a resolution of 30K with charge dependent and the charge state dependent ETD calibration routine options enabled)

*A. baumannii* and Jurkat proteomics samples were analysed using 125-min analytical runs, altering the buffer composition from 3% Buffer B (0.1% formic acid, 77.9% acetonitrile, 2% DMSO) to 28% B over 90 min, then from 28% B to 40% B over 9 min, then from 40% B to 80% B over 3 min. The composition was held at 80% B for 2 min, and then dropped to 3% B over 2 min before being held at 2% B. The Lumos Mass Spectrometer was operated in a data-dependent acquisition mode with datasets collected with either HCD or EThcD fragmentation. For HCD dataset a single Orbitrap MS scan (350–2,000Lm/z, maximal injection time of 118 ms, an Automated Gain Control (AGC) set to a maximum of 100% and a resolution of 60k) was collected every 3 s followed by a Orbitrap MS/MS HCD scans of precursors (NCE 30%, maximal injection time of 80 ms, an AGC set to a maximum of 250%L and a resolution of 30k).For EThcD samples, a single Orbitrap MS scan (450– 2,000Lm/z, maximal injection time of 246 ms, an Automated Gain Control (AGC) set to a maximum of 100% and a resolution of 120k) was collected every 3Ls and Orbitrap MS/MS EThcD scans of precursors (NCE 30%, maximal injection time of 250Lms, an AGC set to a maximum of 500%L and a resolution of 30k), with charge dependent and the charge state dependent ETD calibration routine options enabled.

### Proteomics analysis

Proteomic datasets were searched with Fragpipe (v23.0) (50–56) using their respective proteomes (Avidin: UniProt - P02701; *A. baumannii* D1279779: NCBI: GCA_000186665.4; Human: UniProt – UP000005640). Avidin samples digested with pepsin were searched with non-specific specificity allowing carbamidomethylation of cysteines as a fixed modification and oxidation of methionine on up to 3 positions within peptides. Carboxyl derivatization of glutamic acid/aspartic acid or the protein C-termini was searched as variable modifications on up to 5 positions (AETMA: 84.1054 Da, Girard’s T Reagent: 113.0953 Da or ethanolamine: 43.0422 Da) to assess labelling efficacy, or as fixed modifications to allow the identification of protein C-termini. To improve identification of AETMA, and Girard’s T Reagent modified peptides remainder fragment masses 25.03163 for AETMA and 54.0218, 26.0269 for GT) and neutral losses (-59.07377, -118.14754, -177.22131 for AETMA and for GT) were included as labile mode and localise mass shift switched on. All searches were performed using a protein FDR of 1%. *A. baumannii* samples were searched with a Arg-C specificity due to the blocking of lysine with dimethylation. Ethanolamine and AETMA modifications were searched with remainder/neutral losses as above allowing modification on E/D and the C-terminal modification as a variable modification to assess labelling efficacy and as a fix modification to allow the identification of Protein C-termini. To allow the searching of the human proteome a focused protein database (57, 58) was defined using the “Open” workflow to enable the identification of subset of proteins observable from the proteome. Jurkat cell datasets were then searched as described above for *A. baumannii* samples using the open search defined focused protein database.

The combined_modified_peptide.tsv and combined_ion.tsv was used for determination of efficiency and charge distribution of the peptides. Calculation of labelling efficiency for each unique peptide were undertaken using the combined precursor intensity of all detected unmodified and modified variants with labelling defined as no labelling (unmodified), partially labelled (at least 1 unlabelled Asp/Glu), and complete labelling (all Asp/Glu residues within a peptide labelled) was determined as a percentage, and averaged across all peptides. For charge distribution, only the precursor intensity of the modified peptides was considered. For determination of Protein/peptide C-termini, the resulting “psm.tsv” files from individual replicates were combined within R and assignments with MSfragger Hyperscores > 10. All figures were generated using R with spectral figures annotated using python (59). The combined_protein.tsv and combined_modified_peptide.tsv were used for statistical analysis using Perseus (v.1.6.0.7). Only proteins/peptides which were quantified in at least three replicates in at least one of the groups were considered for statistical analysis. Remaining missing values were imputed based on normal distribution (σ-width = 0.3, σ-downshift = -1.8). Significance was determined using a student’s two-way t-test with a threshold set to Log_2_(fold-change) > |1| and - Log_10_(p-value) > 1.3 (p < 0.05). These were visualised as volcano plots using the ggplot package in R.

### Statistics and Reproducibility

Statistical analyses of biological samples were undertaken on a minimum of three technical replicates for avidin and four biological replicates for biological samples. A biological replicate is defined as a separately grown culture treated with a given induction or growth condition. All mass spectrometry data (RAW files, FragPipe outputs, Rmarkdown scripts, and output tables) have been deposited into the PRIDE ProteomeXchange repository (60, 61). All PRIDE accession numbers and descriptions of the associated experiment are provided within Supplementary Table 1.

## Results and discussion

### On-bead carboxylic derivatization allows C-terminal peptide labelling

To explore the potential of on-bead C-terminomics, we first assessed the efficiency of carboxyl derivatization using two previously reported C-terminal protein labelling reagents: AETMA and ethanolamine (30, 32). Using avidin as a model protein, PAC/SP3-based amine group protection via dimethylation was undertaken according to Weng et al. (39). Carboxyl amidation was subsequently performed using EDC/HOBt coupling with DIPEA in a single overnight labelling round as described by Solis et al. (30). Pepsin-based digestion of avidin confirmed successful derivatization of carboxyl groups, as assessed by modification of Asp/Glu residues, yet revealed incomplete carboxyl labelling (Table S2). Quantification of carboxyl labelling revealed that 32 ± 8.6% of identified peptides were fully labelled with ethanolamine, while 49 ± 7.3% of peptides were completely labelled with AETMA (Figure 1A). Consistent with this, intact protein analysis supports the labelling of 45% and 60% of available sites (12 potential sites) with ethanolamine and AETMA, respectively (Figure S1A-B). As previously reported, the derivatization of carboxyl groups resulted in a notable shift in the observed charge states of ethanolamine-labelled and AETMA-labelled peptides compared to dimethylated peptides (Figure 1B). Importantly, both ethanolamine and AETMA successfully resulted in C-terminal labelling of avidin (Figure 1C-D), with manual inspection also revealing the presence of a dominant neutral loss of 59.074 Da within AETMA-modified peptides upon collision activation (Figure 1C), consistent with the loss of trimethylamine previously reported from AETMA-labelled oligosaccharides (62). Taken together, these results support that PAC/SP3 allows on-bead derivatization, yet revealed modest overall efficiency using in-solution carboxyl labelling protocols.

**Figure 1.**
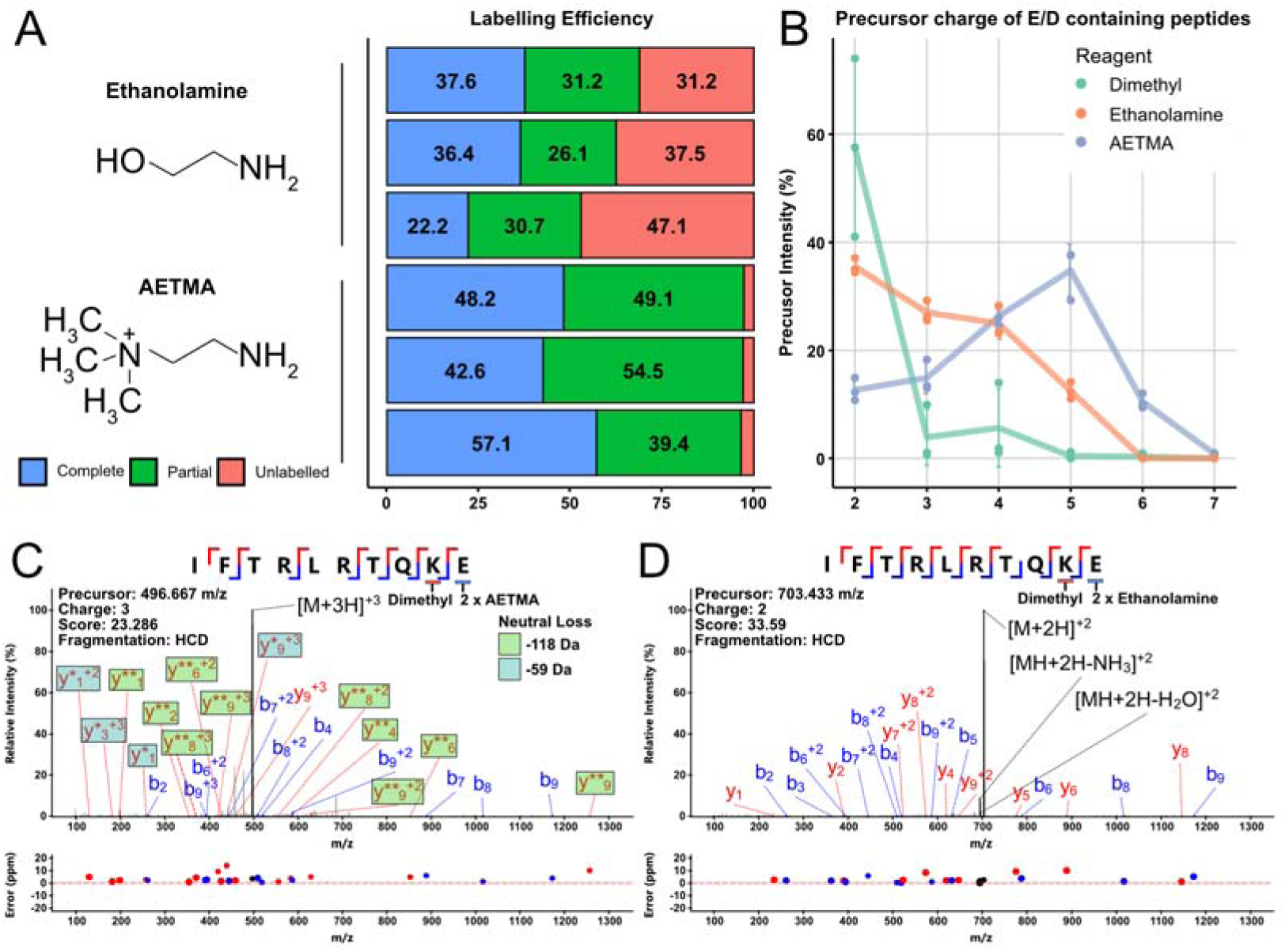
Carboxyl derivatization of Avidin on SP3 beads. **(A)** Carboxyl on beads Ethanolamine and AETMA derivatization using in solution optimised protocols results in incomplete labelling. The precursor intensity (%) of relative labelling of the two reagents was either complete (all sites occupied), partial (at least 1 site occupied), and unlabelled (site/s present but no labelling observed). Individual data points are three technical replicates where labelling was performed in separate reactions. **(B)** The % precursor intensity of charged peptides that contain E/D or labelled C-termini for each of the amine reagents. Individual data points are three technical replicates where labelling was performed in separate reactions. An example of **(C)** HCD and **(D)** EThcD spectra of the Avidin protein C-terminal peptide ^119^IFTRLRTQKE(84.1054)c(84.1054)^128^. Labile loss on y-ions is indicated by * where each * represents -59.07 Da.

### Sequential on-bead labelling improves carboxylic derivatization efficiency

To enable robust C-terminomics, near-complete carboxylic labelling is essential. To improve efficiency, we reasoned that the addition of fresh labelling reagents and extended labelling duration may improve the overall labelling completeness. To assess this, we undertook sequential labelling rounds over 48 hours, where freshly prepared labelling reagents were added after 24 hours. To extend our assessment of carboxyl labelling reagents, we also assessed the efficiency of sequential labelling with an additional carboxyl labelling regent, Girard’s T Reagent. Compared to single labelling, sequential labelling increased the completeness of AETMA labelling to 85.5 ± 2%, while labelling remained relatively low for ethanolamine at 41 ± 4.5% (Figure 2A, Table S3). In line with the labelling of AETMA, Girard’s T reagent resulted in 68 ± 2.3% labelling efficiency. Consistent with our single labelling results, sequential labelling increased the observed charge states of identified peptides, with both AETMA and Girard’s T reagent resulting in similar increases in observed charge states (Figure 2B). Like AETMA, Girard’s T Reagent-derivatized peptides also resulted in the generation of neutral loss species under collisional activation (-59.074 and -87.068 Da) (Figure S2). While the benefits of fixed-charge derivatizations for enhancing electron-based dissociation of peptides have been previously explored (16, 18, 32), we examined the impact of C-terminal protein labelling on ETD fragmentation complementarity. Examination of the lowest observed charge state of the C-terminal avidin peptide ^120^FTRLRTQKE^128^, with each labelling reagent, revealed enhanced z/c ion generation in fixed-charge-derivatized peptides (Figure 3A-B), consistent with previous reports (18). Taken together, the use of sequential on-bead labelling improves carboxyl derivatization, allowing effective C-terminal labelling with the fixed-charge reagents AETMA or Girard’s T reagent resulting in increased peptide charge states and richness of ETD spectra.

**Figure 2.**
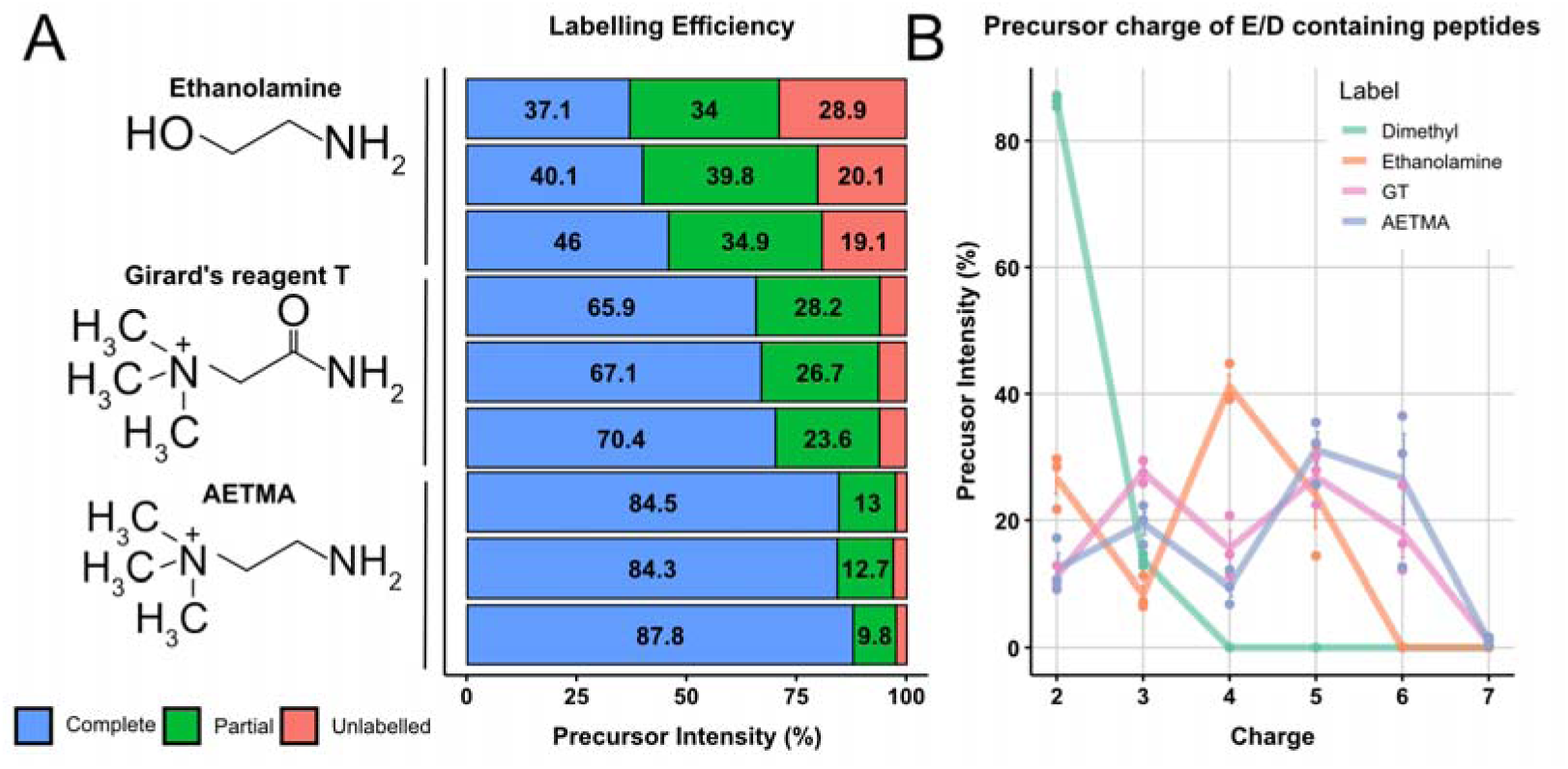
Improving labelling efficiency. To improve labelling efficiency, an additional day of carboxyl group labelling was performed on intact avidin using labelling reagents Ethanolamine, AETMA and Girard’s reagent T. **(A)** The precursor intensity (%) of relative labelling of the three reagents as either complete (all sites occupied), partial (at least 1 site occupied), and unlabelled (site/s present but no labelling observed). **(B)** The % precursor intensity of charged peptides that contain E/D or labelled C-termini for each of the amine reagents. Individual data points are three technical replicates where labelling was performed in separate reactions.

**Figure 3.**
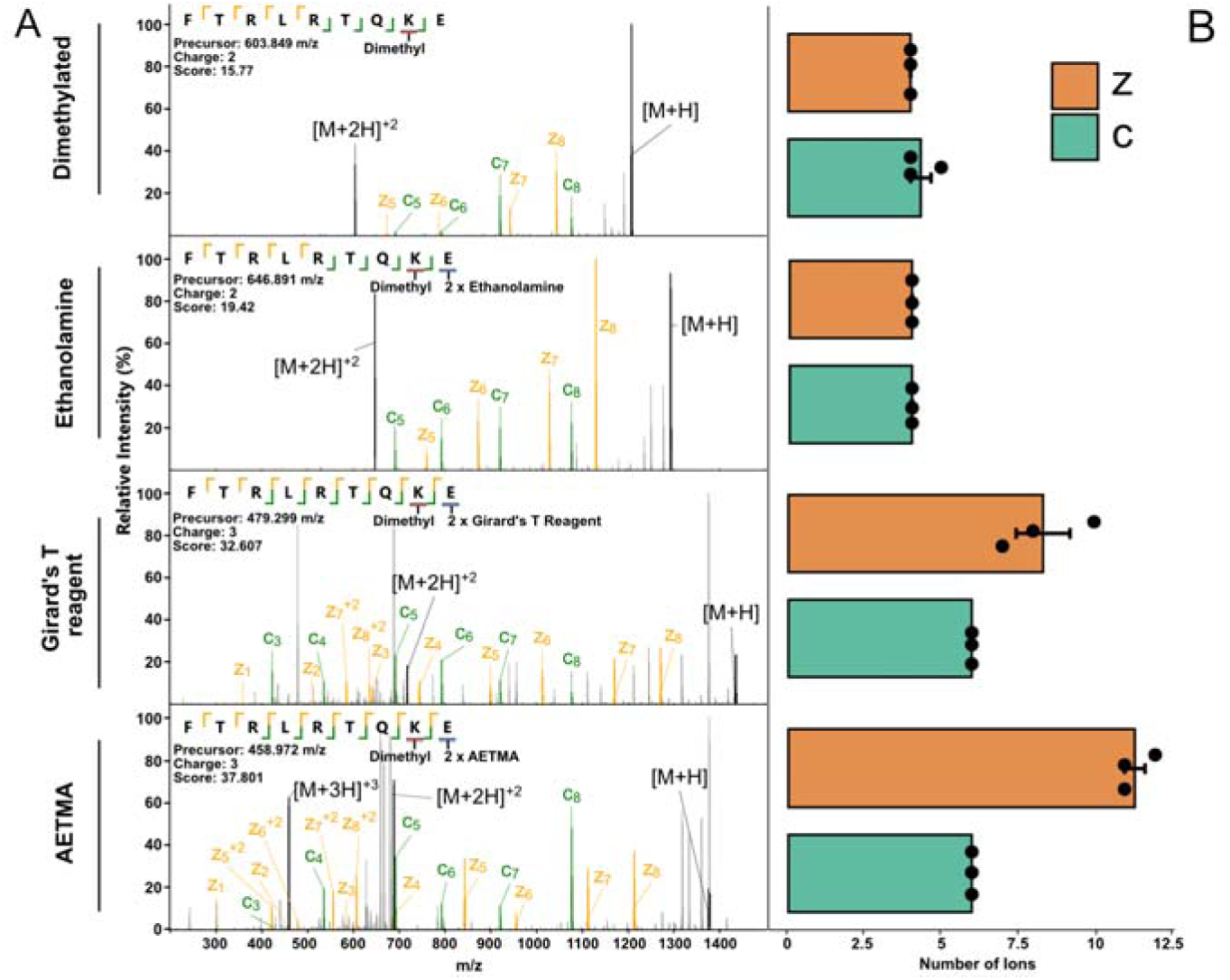
Improved ETD fragmentation of protein C-terminal avidin peptide. **(A)** EThcD spectra of the protein C-termini peptide of avidin, ^120^FTRLRTQKE^128^, where both the aspartic acid and C-termini are labelled with either ethanolamine, Girard’s T Reagent or AETMA showing improved fragmentation. **(B)** Number of detected z/c ions within a 10-ppm threshold detected across spectra in 3 technical replicates for each of the reagents, with Girard’s T reagent and AETMA having the largest increase in z-ions.

### On-bead carboxyl derivatization allows the identification of C-terminal peptides at a proteome scale

To assess the capacity of on-bead carboxyl labelling for proteome-scale identification of protein C-termini, we utilized whole-cell lysates of *A. baumannii* D1279779 (63) as a model system. Using sequential labelling, we derivatized *A. baumannii* samples with AETMA or ethanolamine and assessed the identification of C-termini using both collision (HCD) and electron (EThcD) fragmentation. At a proteome level, AETMA and ethanolamine labelling resulted in an overall completeness of 56.5 ± 1.76 % and 29 ± 2.5 % respectively (Figure S3A, Table S4). Consistent with our single-protein labelling, AETMA derivatization resulted in higher labelling efficiency yet only achieved modest labelling completeness. Interestingly, examination of the overall identification rate of AETMA-derived peptides revealed reduced performance compared to ethanolamine for both electron and collision-based fragmentation (Figure 4A-B). Examination of the AETMA-labelled peptide hyperscores revealed that the use of collision-based fragmentation resulted in overall lower scoring identifications and a negative correlation between scores and number of labelling events (Figure S3D) compared to ethanolamine (Figure S3B). This trend was not observed with ETD fragmentation of AETMA-labelled peptides (Figure S3C, E), but a reduction in the total number of identified AETMA peptides was still observed (Figure S4A-B). Combined these observations support that AETMA labelling is associated with reduced identification rates using both collision and electron-based fragmentation, despite higher labelling efficiency.

**Figure 4.**
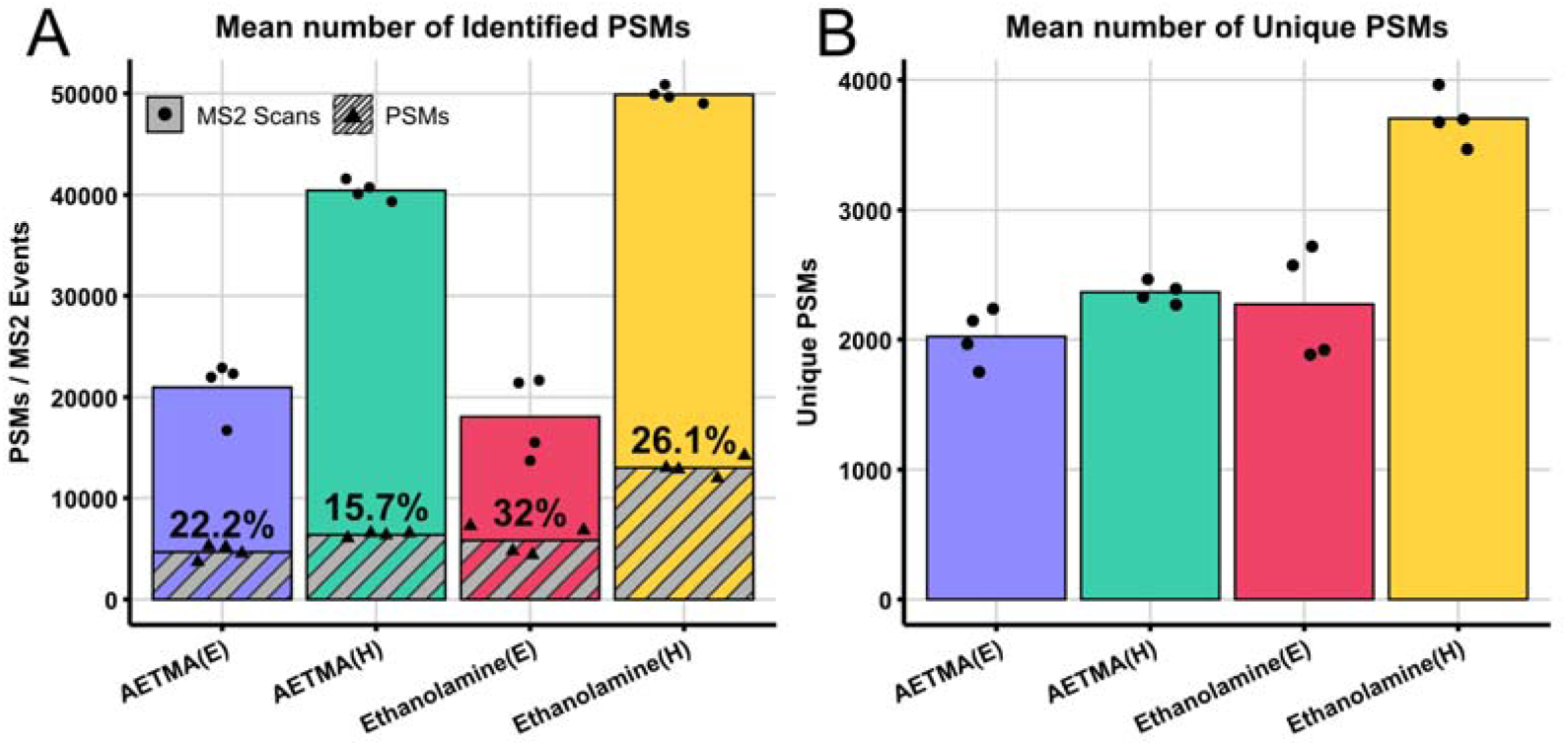
Identification rate and unique PSMs identified within *A. baumanni* D1279779 following on-bead carboxyl derivatization. **(A)** The mean number of MS2 events across the replicates for each MS run, and mean number of peptide spectrum matches (PSMs). **(B)** The total number of unique PSMs identified across MS run.

Across these two reagents we observed a total of 153 protein C-terminal peptides, corresponding to 137 unique proteins (Figure 5A, Figure S4A, Table S4), these included 37 and 55 unique C-termini identified with AETMA and ethanolamine, respectively; 61 were identified across both reagents (Figure 5A, Table S4). In addition to C-termini, this method also allows for identification of N-termini with a total of 297 unique protein N-terminal peptides detected, corresponding to 269 unique proteins, and in total 7,275 peptides were identified corresponding to 1,142 unique proteins (Figure S4B-C). Examination of the amino acid composition of identified C-terminal peptides revealed notable differences in the total number of positively charged residues (H/K/R) observed with AETMA compared to ethanolamine labelling (Figure 5B). Specifically, of the 37 C-terminal peptides uniquely identified with AETMA, 64.9% (Figure 5B) lacked positively charged residues (Figure S5A-B). For example, the peptide ^763^TYGLSYFFNY^772^ derived from the protein C-terminus of AGH35524.1 was only identified with AETMA derivatization (Figure 5E-F). In contrast, for the 55 C-terminal peptides uniquely observed with ethanolamine labelling, 92.7% (Figure 5B) possessed internal basic residues (Figure S5C-D). Consistent with these trends, motif sequence analysis revealed differences in the frequency of basic residues observed within AETMA-labelled and ethanolamine-labelled protein C-terminal peptides (Figure 5C-D). These findings demonstrate that on-bead carboxylic labelling allows the identification of C-terminal-labelled peptides from both AETMA/ethanolamine labelling. While many C-terminal peptides can be identified with either labelling reagents we find AETMA uniquely enables the identification of C-termini lacking internal basic residues.

**Figure 5.**
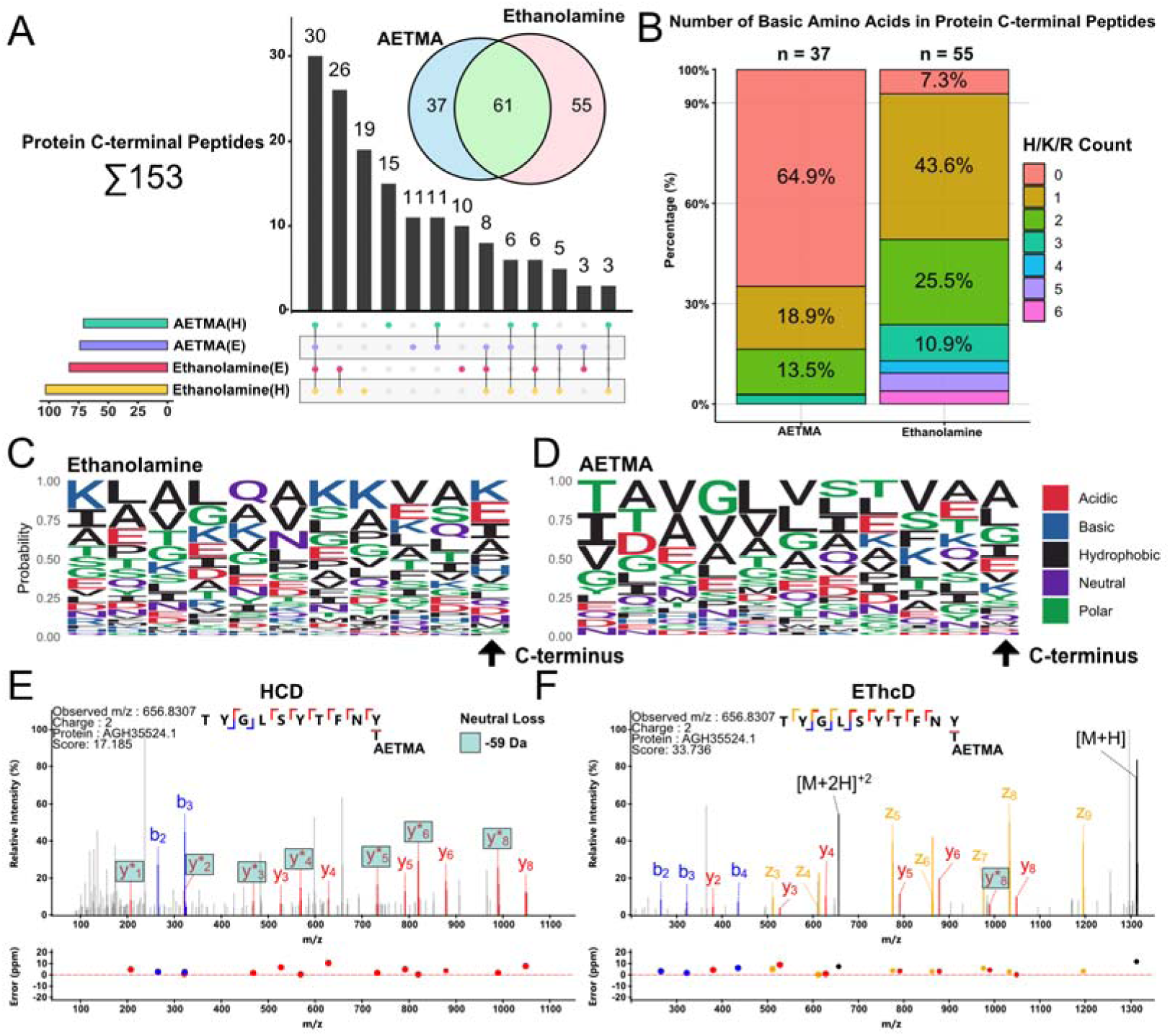
AETMA and ethanolamine identify unique C-termini within *A. baumannii* D1279779. Mass spectrometry analysis was performed on *A. baumannii* D1279779 proteins labelled with AETMA or ethanolamine with HCD (H) or EThcD (E) fragmentation. **(A)** Upset plot of unique and shared Protein C-terminal peptides between AETMA and ethanolamine labelled samples. **(B)** The number of basic amino acids present in PSMs for the two different labels. **(C)** The amino acid frequency of protein C-terminal peptides measured as a percentage difference of AETMA compared to ethanolamine labelled peptides. Positive values indicate an enrichment of amino acids in AETMA, and depletion for negative values, and vice-versa for ethanolamine. **(D)** Peptide sequence motif of unique protein C-terminal peptides for AETMA and ethanolamine with the final position being the C-terminus of the protein. Protein C-terminal peptides are considered based on genomic ORF. **(E-F)** HCD and EThcD spectra of the C-terminal peptide ^763^TYGLSYFFNY^772^ of AGH35524.1 labelled with AETMA, with * indicates neutral loss of 59 Da.

### On-bead derivatization allows proof of concept for combined N- / C-terminomics in eukaryotic protein samples

The ability to identify C-terminal peptides from on-bead derivatization supports that single pot N-/C-terminomics may be achievable. To explore this within a eukaryotic context, we utilized etoposide-induced apoptosis of Jurkat cells, a widely used model within previous degradomics studies (64–66). Confirming induction of apoptosis following etoposide treatment using the caspase activity-based probe LE22 (46) (Figure S6), we applied N- and C-terminal derivatization, employing collision (HCD) and electron (EThcD) based fragmentation on samples labelled with AETMA or ethanolamine as above (Figure S7A). Across samples, we identified 1,334 proteins, with 628 proteins identified across all labelling/fragmentation conditions (Figure S7B; Table S5). A total of 7,016 peptides were identified with 1,476 peptides observed across all labelling/fragmentation conditions (Figure S7C; Table S6). Assessment of these peptides demonstrated AETMA labelling provided a higher proportion of labelled D/E residues compared to ethanolamine, regardless of fragmentation method (72-78% versus 62-65%) (Figure S8A), as well as the identification of high (>+3) charge state peptides (Figure S8B-C). Additionally, of the total peptides identified, 13-22.5% (1,722 unique peptides across all conditions) contained dimethylated N-termini (Figure S8D), in line with the 17.31% of peptides previously reported using PAC/SP3-enabled N-terminomics (42). For C-termini, only 0.8-1.51% (153 unique peptides across all conditions) were identified, with the highest proportion reported following AETMA labelling and EThcD fragmentation (1.51%, 53 C-termini) (Figure S8D). Together, these data suggest AETMA offers better carboxyl labelling efficiency and increases the charge state of labelled peptides

Of the 153 C-termini identified, only 23 C-termini were shared between AETMA and ethanolamine labelling (Figure 6A). Across the unique C-termini, the majority were identified using a single labelling/fragmentation strategy (Figure 6A), which is opposite to what is observed in the *A. baumannii* lysates (Figure 4A) and may be due to the increased complexity of the eukaryotic proteome. Consistent with the analysis of *A. baumannii*, AETMA labelling resulted in the identification of C-terminal peptides lacking basic residues (H/K/R) (Figure 6B), such as the C-terminal peptide ^97^TLYGFGG^103^ of Histone H4 (P62805) identified following AETMA labelling and EThcD (Figure 6C). By comparison, all detected C-termini labelled with ethanolamine contained ≥1 charged residue (Figure 6B). This was further supported by the fact that ethanolamine-labelled C-terminal peptides contained a higher proportion of lysine residues compared to AETMA-labelled peptides, which contained more glycine and alanine residues (Figure S9). Among the 153 identified C-terminal peptides, no P1 aspartic acid cleavage events were observed, suggesting that while C-terminal-labelled peptides can be identified within a eukaryotic context, the low identification frequency of these peptides may limit C-terminomics without enrichment. Additionally, no P1 glutamic acid cleavage events were detected (Figure S9A-B). Therefore, the labelling of these acidic residues may be limited by steric hinderance due to the double labelling event on the side chain and C-terminus. Despite the lack of detectable C-terminal caspase events, N-terminal caspase cleavage events were readily observed, including several previously reported caspase cleavage events (Figure 7): vimentin (VIM) at D^85^↓F^86^, D^257^↓V^258^, and D^259^↓V^260^; polypyrimidine tract-binding protein 1 (PTBP1) at D^172^↓A^173^; and far upstream element-binding protein 2 (KHSRP) at D^102^↓F^103^ (66). Combined, these results demonstrate that on-bead labelling of carboxyl groups does not impede N-terminomic analysis of unenriched samples. However, the modest frequency of C-terminal identification limits the utility of this approach in its current form, supporting the need for further refinement.

**Figure 6.**
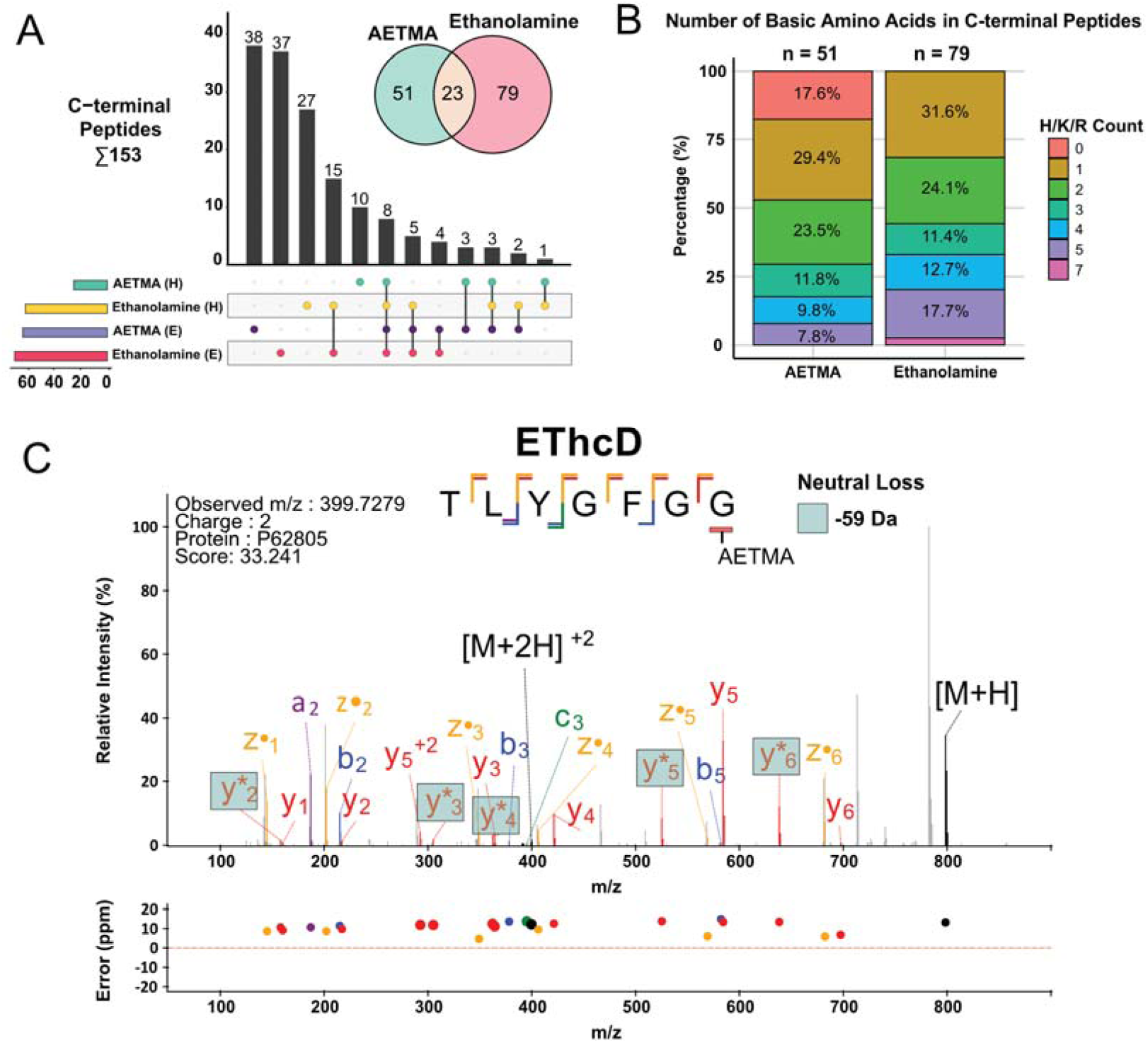
C-terminal labelling with AETMA or Ethanolamine provide complementary identifications in apoptotic Jurkat T-lymphocytes. Apoptosis was induced in human Jurkat cells by administrating etoposide for 8 hours, with DMSO as a control. Protein lysates underwent carboxyl labelling with either AETMA or ethanolamine prior to mass spectrometry analysis with HCD (H) or EThcD (E). fragmentation. **(A)** Upset plot and Venn diagram of unique and shared C-termini between AETMA- and ethanolamine-labelled samples upon differential fragmentation methods. **(B)** The number of basic amino acids present observed in PSMs for the two different labels. **(C)** EThcD spectra for a Protein C-terminal peptide, ^97^TLYGFGG^103^ (P62805) unique to AETMA samples lacking H/K/R charged residues.

**Figure 7.**
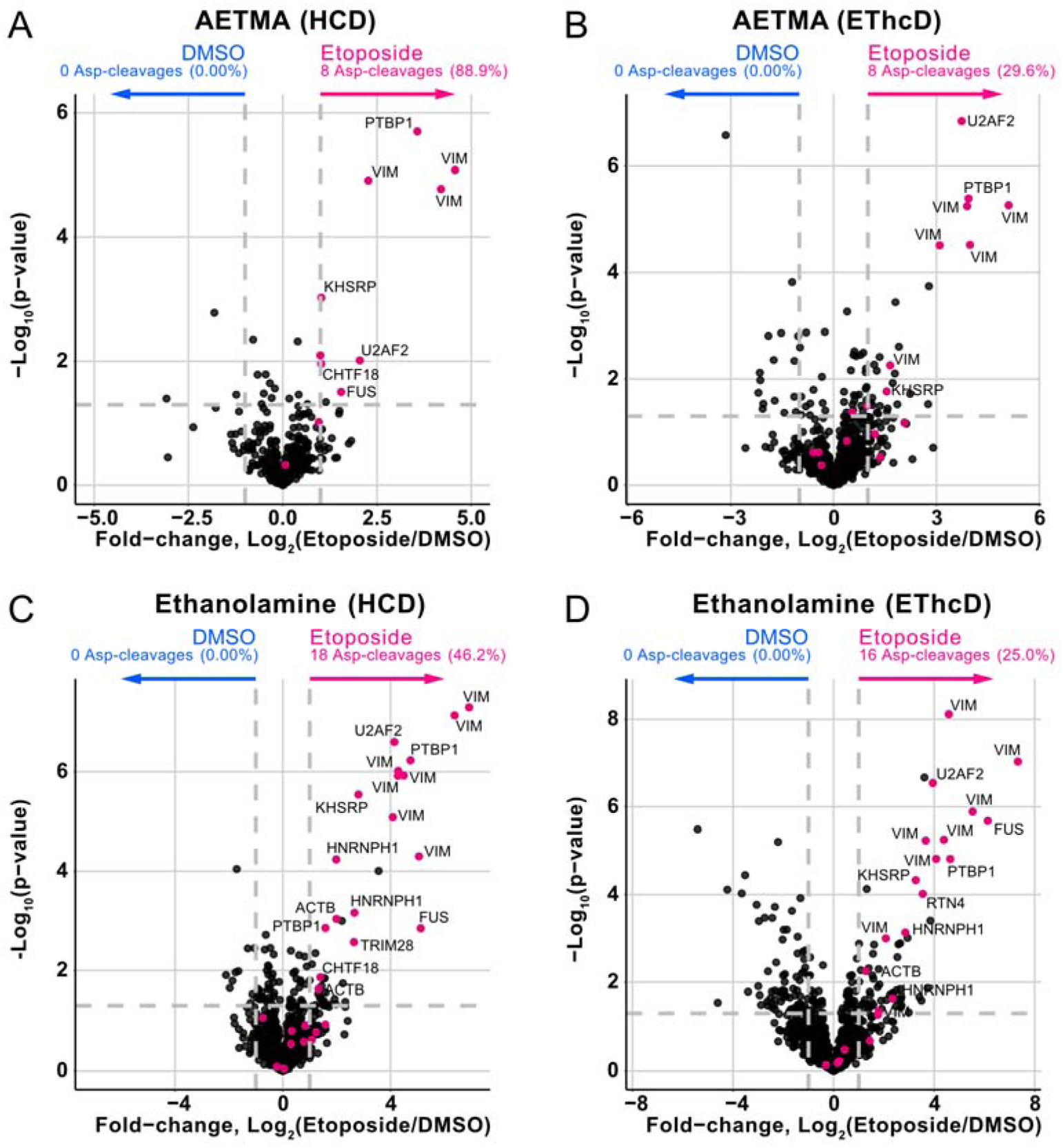
Caspase-dependent cleavage events identified following etoposide-induced apoptosis. **(A-D)** Peptides were filtered to contain only those labelled by dimethylation at the N-terminus, representing native and neo-N-termini. Volcano plots of N-termini changes between apoptotic and control (DMSO) Jurkat cells (n = 4/group). A student’s two-way t-test was used to determine significance at |Log_2_(Etoposide/DMSO) > 1| and -Log_10_(p-value) > 1.3 (p < 0.05). P1 aspartic acid cleavages indicative of pro-apoptotic caspase activity are highlighted in pink.

## Conclusion

Within this study, we have assessed the capacity of on-bead carboxyl labelling to allow the identification of C-terminal peptides. We show that PAC/SP3-based labelling with the carboxyl labels AETMA, Girard’s T Reagent, or ethanolamine allows single-pot labelling following on-bead dimethylation. By extending carboxyl group labelling, modest (60-90%) labelling efficiency using AETMA or Girard’s T Reagent, but not ethanolamine can be achieved. Nevertheless, ethanolamine labelling in both prokaryotic and eukaryotic contexts resulted in overall higher numbers of unique C-terminal peptide identifications. Our observation of abundant neutral losses within AETMA-labelled peptides following collision-based fragmentation suggests that the resulting higher spectral complexity may be the driver of these observations, with similar results observed when using Girard’s T Reagent. Overall, AETMA and ethanolamine displayed complementarity in the C-terminal peptides identifiable with these labelling agents’ support both their utility for C-terminomic analysis.

To date, there has been limited development in techniques for simultaneous assessments of N- and C-termini within proteomic samples. A key challenge for the field has been how to achieve N/C-termini labelling while maintaining compatibility with downstream preparation steps. As reagents required to achieve C-terminal labelling, such as EDC, are unstable in aqueous solutions (67), by leveraging PAC/SP3 we show this is an ideal platform for proteomic labelling under conditions typically poorly compatible with protein samples. Whilst previous methods utilise filter-based buffer displacement for consecutive chemical labelling steps (29, 30, 68, 69), the ability to achieve labelling in a single pot and undertake washes in a touchless manner both simplifies labelling as well as reduces sample loss. Additionally, the ability for PAC/SP3 platforms to be readily automated (41) supports this platform could be used to improve the throughput of N/C-termini degradomics workflows. While this work demonstrates that combined N- and C-terminome information can be achieved in a single workflow as a proof of concept, the current efficiency of C-labelling suggests this is most suitable for low-complexity samples, such as recombinant proteins. This work further demonstrates the versatility of PAC/SP3 sample preparation approaches and the need for refinement of carboxyl labelling chemistry to enable more complete coverage of the C-terminome.

## Data availability

All raw files, Fragpipe output files, FASTA files, and the experimental template have been uploaded to PRIDE under accession numbers; PXD068617, PXD068652, PXD068858, PDX068110

## Supporting information

- Supplementary documentation containing; (Table S1) description of the PRIDE proteomeXchange accession numbers for each experiment, (Figure S1) spectra for intact labelling of Avidin, (Figure S2) spectra for AETMA and Girard’s T Reagent labile loss following collision-induced dissociation, (Figure S3) labelling efficiencies and hyperscores of peptide-spectrum matches in *A. baumannii* samples, (Figure S4) overlap of protein and peptide identifications between AETMA and ethanolamine-labelled *A. baumannii* samples, (Figure S5) spectra of C-terminal peptides detected in *A. baumannii* samples, (Figure S6) verification of etoposide-induced apoptosis in Jurkat cells, (Figure S7) overlap of protein and peptide identifications between AETMA and ethanolamine-labelled Jurkat samples, (Figure S8) proportion of peptides detected in Jurkat samples, (Figure S9) C-terminal labelling specificities for Jurkat samples (PDF)
- (Table S2) Avidin peptides identified following a single carboxyl labelling step (XLSX)
- (Table S3) Avidin peptides identified following two carboxyl labelling steps (XLSX)
- (Table S4) Highest scoring unique peptides identified following AETMA and ethanolamine labelling of *A. baumannii* lysates (XLSX)
- (Table S5) Proteins identified following AETMA and ethanolamine labelling of Jurkat cell lysates (XLSX)
- (Table S6) Peptides identified following AETMA and ethanolamine labelling of Jurkat cell lysates (XLSX)

## Acknowledgements

We thank the Melbourne Mass Spectrometry and Proteomics Facility of The Bio21 Molecular Science and Biotechnology Institute for access to MS instrumentation. We thank Dan Polasky for their help and guidance on improving search parameters for searches in MSFragger. N.E.S and K.I.K are supported by an Australian Research Council (ARC) Future Fellowship (FT200100270), an ARC Discovery Project Grant (DP210100362) and a National Health and Medical Research Council Ideas grant (2018980). L.E.E-M. was supported by a National Health and Medical Research Council Ideas Grant (GNT2011119). A.R.Z. was supported by RTP Scholarships from the Australian Government.

## Author Contributions

K.I.K. – Data curation; Formal analysis; Investigation; Methodology; Validation; Visualization; Writing – original draft; Writing – review & editing

A.R.Z. – Data curation; Formal analysis; Investigation; Methodology; Validation; Visualization; Writing – original draft; Writing – review & editing

L.E.E-M. – Funding acquisition; Methodology; Project administration; Resources; Supervision; Writing – review & editing

N.E.S. – Conceptualization; Funding acquisition; Methodology; Project administration; Resources; Supervision; Writing – review & editing

