## Supplementary document for "PAC/SP3 on-bead carboxyl derivatization allows combined C- and N-terminomics"

**Table of contents**

| **Supplementary Tables and Figures** | **Page** |
| --- | --- |
| **Table S1:** Pride Proteomic datasets | 3 |
| **Figure S1:** Intact protein analysis of AETMA and ethanolamine labelled Avidin | 4 |
| **Figure S2:** Labile loss of AETMA and Girard’s T Reagent during HCD fragmentation | 5 |
| **Figure S3:** Number of identified PSMs in *A. baumanni* D1279779 and hyperscore for different number of labels | 6 |
| **Figure S4:** Shared C-/N- and Total Peptides and Proteins of *A. baumannii* D1279779 samples | 7 |
| **Figure S5:** Examples of protein C-terminal peptides from *A. baumanni* D1279779 observed only in AETMA or ethanolamine examples | 8 |
| **Figure S6:** Induction of apoptosis by etoposide in Jurkat cells | 9 |
| **Figure S7:** Overlap of protein and peptide identifications across different labelling and fragmentation strategies | 10 |
| **Figure S8:** AETMA increases carboxyl labelling efficiency and precursor charge states | 11 |
| **Figure S9:** Amino acid preferences for detected C-termini in Jurkat cells | 12 |

**Additional Supplementary Tables**

**Table S2:** Avidin peptide digests single labelling

**Table S3:** Avidin peptide digests double labelling

**Table S4:** *A. baumannii* D1279779 highest scoring unique peptides

**Table S5:** Protein abundance differences between DMSO and etoposide-treated Jurkat cells

**Table S6:** Peptide differences between DMSO and etoposide-treated Jurkat cells

**Supplementary Table 1. Pride Proteomic datasets.**

| Pride accession number (Review login details) | MS instrument | Number of Biological groups, replicates and total datafiles | Description of dataset |
| --- | --- | --- | --- |
| PXD068617  Username:  Password: Q2KTbbDMK4qJ | LUMOS | 4 groups each with 3 replicates (two search types) total data files is 24. Version 1 is HCD/ETD and Version 2 is HCD/EThcD | Pepsin digest of the protein analysis to assess carboxyl derivatization |
| PXD068652  Username:  Password: nT3flrelLayq | LUMOS | 4 groups each with 4 replicates total data files is 16 | Pepsin digest of dimethylated, ethanolamine, AETMA and Girard’s T reagent samples with extended labelling |
| PXD068858 Username: Password: Ao63a3NUfktJ | LUMOS | 2 groups each with 4 replicates (two search types) total data files is 16. One set of 8 were fragmented with HCD, and other set were fragmented with EThcD | Trypsin digest of *A. baumanni* D1279779 samples labelled with either AETMA or ethanolamine |
| PXD068110  Username:  Password: 0kth0jvGJtms | LUMOS | 2 groups each with 4 replicates, split into 4 separate experiments based on labelling reagent and fragmentation method. Total data files is 32 | Apoptotic Jurkat cell proteomic and degradomics analysis |

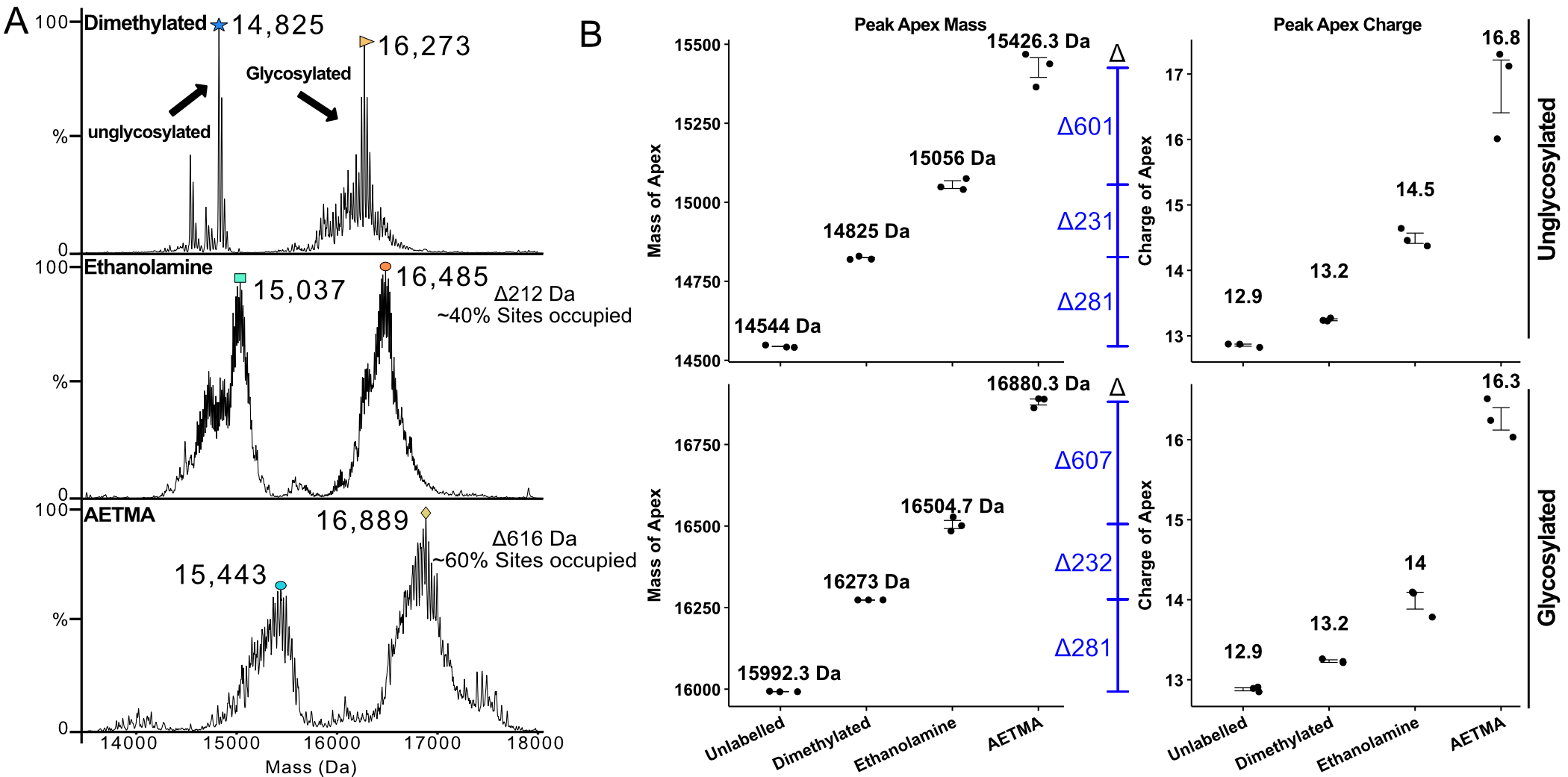

**Figure S1. Intact protein analysis of AETMA and ethanolamine labelled Avidin.** Intact protein analysis of unlabelled (reduced/alkylated), demethylated, ethanolamine and AETMA labelled avidin. **(A)** Representative mass spectra of a single replicate of dimethyl-, ethanolamine- or AETMA-labelled avidin depicting both unglycosylated and glycosylated variants. **(B)** Mass of individual replicates based on the peak apex for both the unglycosylated and glycosylated avidin peaks, and the average charge of those peaks. All data were collected on a Waters QTof, then deconvoluted using UniDec. Individual technical replicates represent separate reactions.

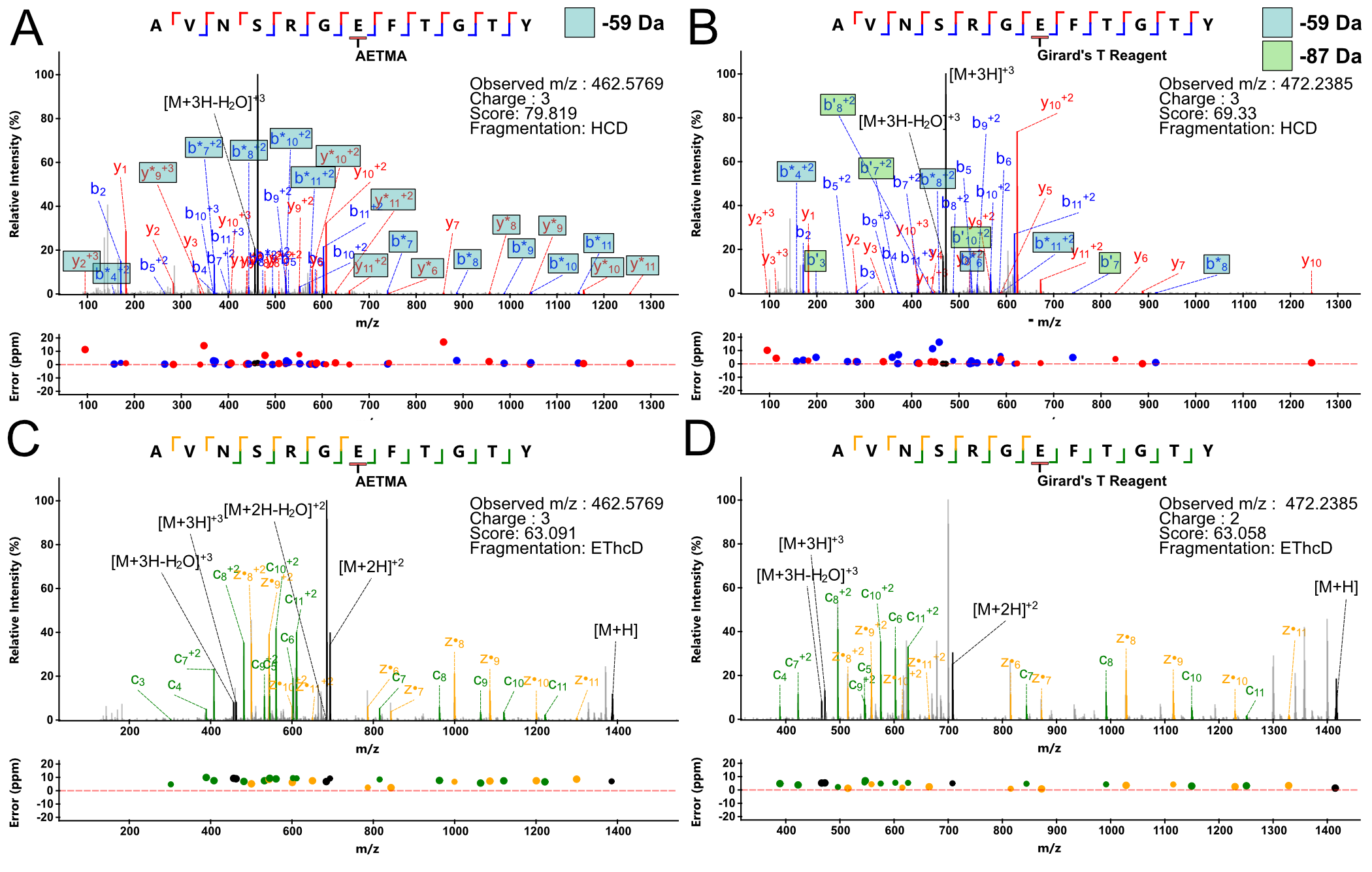

**Figure S2. Labile loss of AETMA and Girard’s T Reagent during HCD fragmentation.** Examplary spectra depicting neutral loss occurring during HCD fragmentation for AETMA and Girard’s T Reagent. **(A-B)** HCD spectra of Avidin peptide ^22^AVNSRG**E**FTGTY^33^ derivatised at single glutamic acid side for AETMA and Girard’s T Reagent. b/y ions with neutral loss of 59 Da are denoted with *; ’ denotes -87 Da loss. **(C-D)** EThcD spectra for the AVNSRG**E**FTGTY peptide with c/z● annotated.

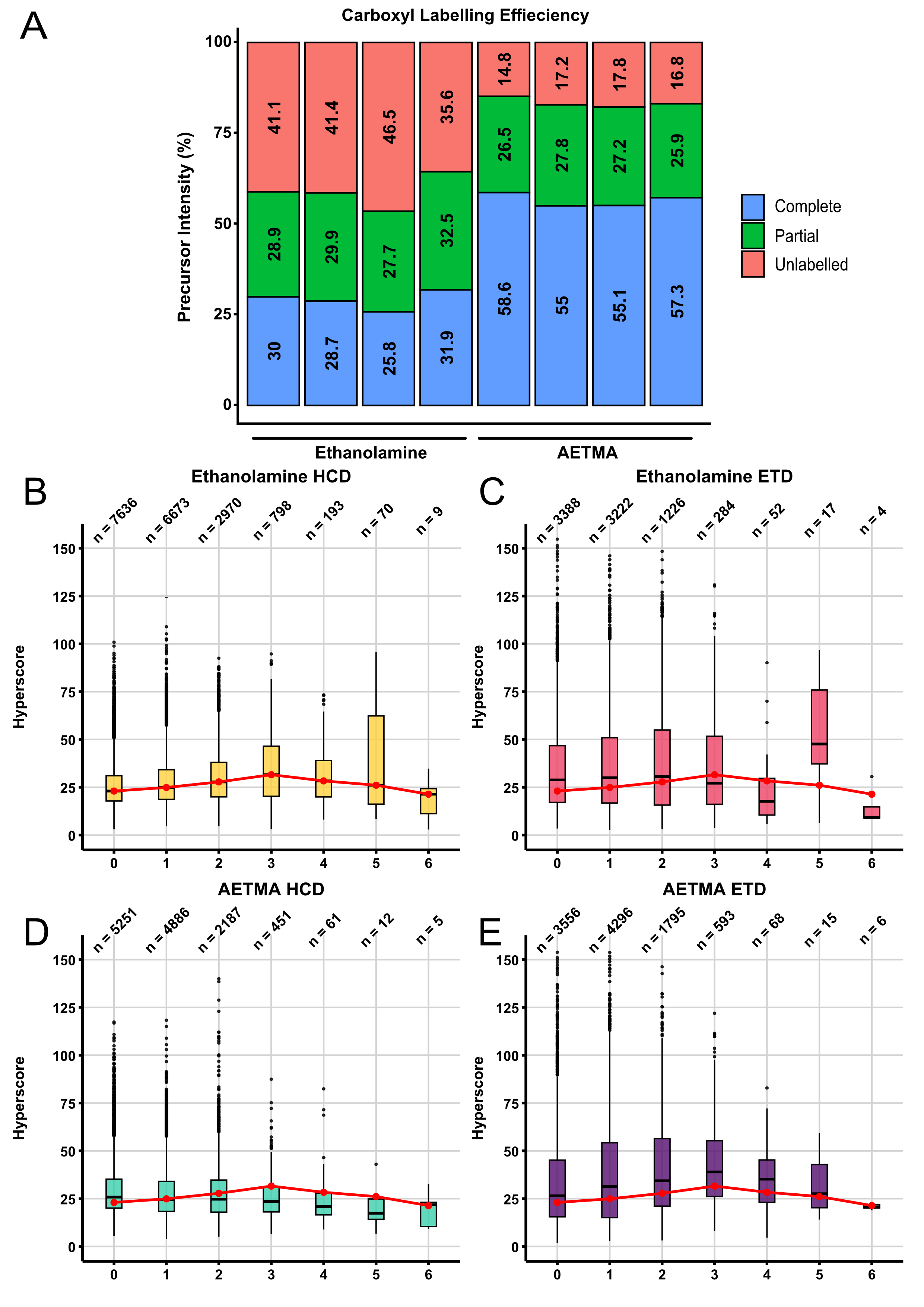

**Figure S3. Number of identified PSMs in *A. baumanni* D1279779 and hyperscore for different number of labels.** **(A)** The precursor intensity (%) of relative labelling of the two reagents as either complete (all sites occupied), partial (at least 1 site occupied), and unlabelled (site/s present but no labelling observed). Individual data points represent four technical replicates . **(B-E)** The hyperscore of peptides containing 0 to 6 labels (AETMA or Ethanolamine) for each of the MS runs, where n is the number of PSMs.

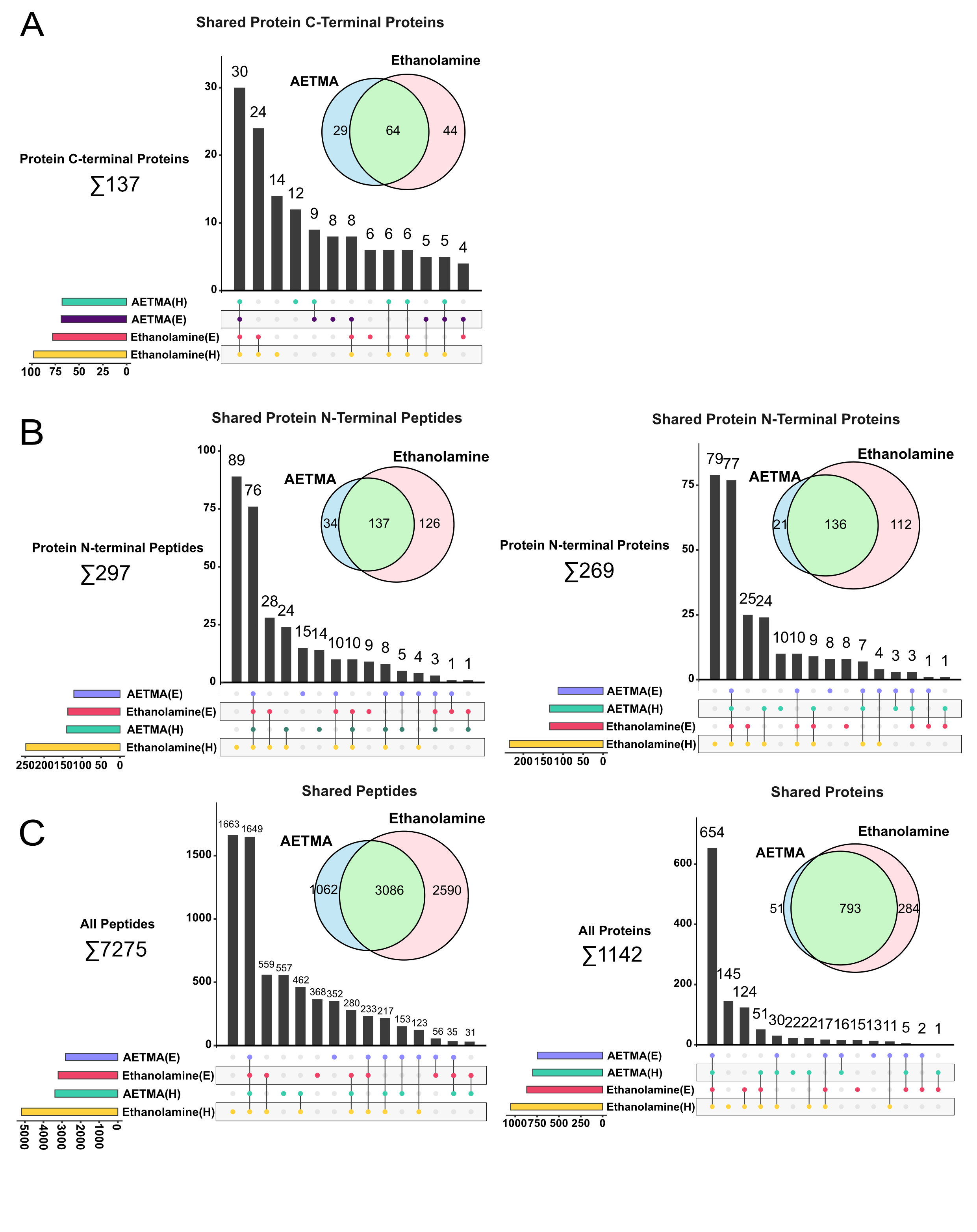

**Figure S4. Shared C-/N- and Total Peptides and Proteins of *A. baumannii* D1279779 samples. (A)** Upset plot of unique and shared Protein C-termini peptides between AETMA and ethanolamine labelled samples. **(B)** Upset plot of unique and shared Protein N-termini Proteins and peptides between AETMA and ethanolamine labelled samples. **(C)** Upset plot of unique and shared Protein and peptides between AETMA and ethanolamine labelled samples.

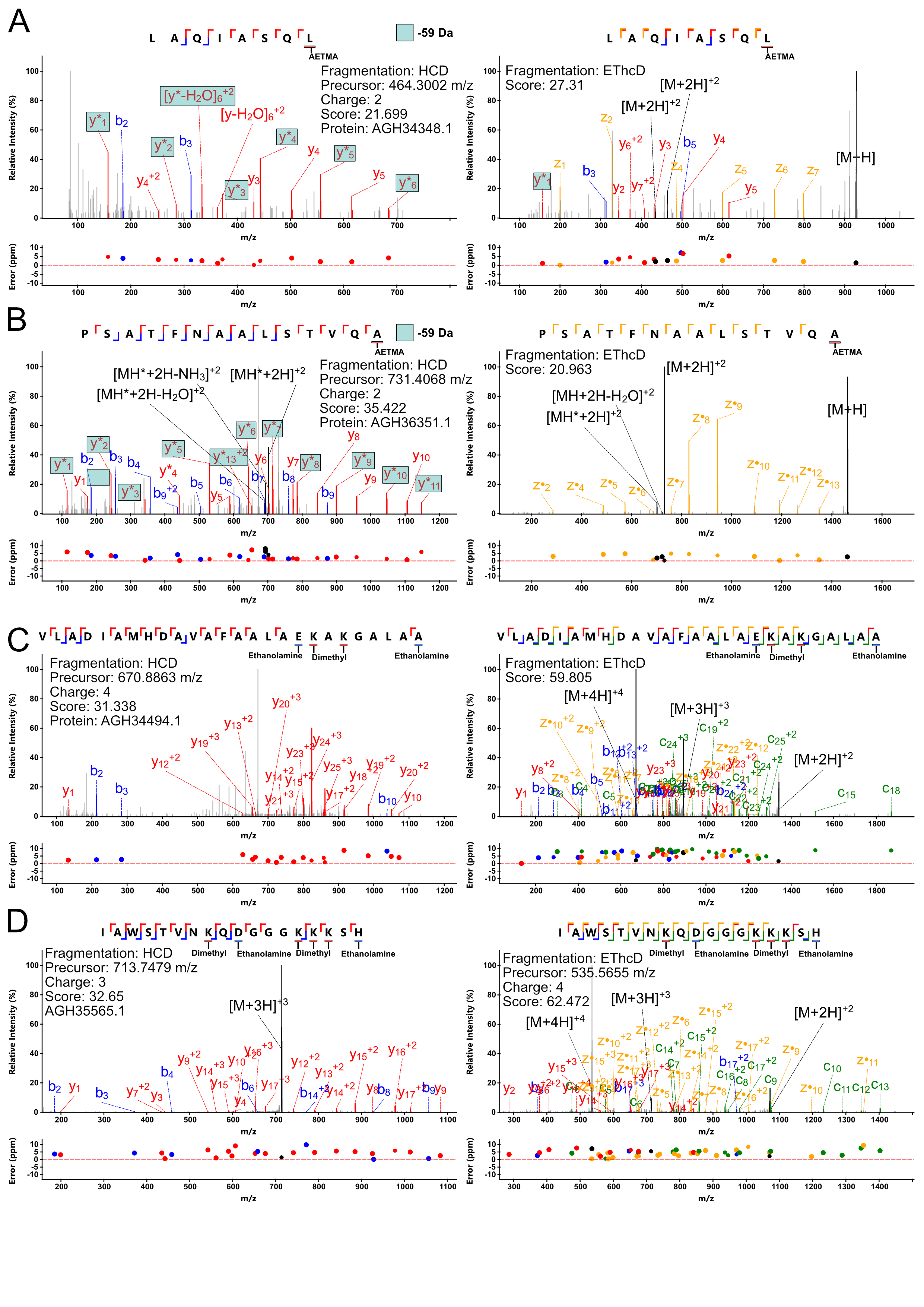

**Figure S5. Examples of protein C-terminal peptides from *A. baumanni* D1279779 observed only in AETMA or ethanolamine examples.** Exemplary HCD and EThcD of protein C-terminal peptides ^199^LAQIASQL^206^ (AGH34348.1) and ^732^PSATFNAALSTVQA^745^ (AGH36351.1) only observed in AETMA **(A-B)**, or ^94^VLADIAMHDAVAFAALAEKAKGALAA^119^ (AGH34494.1) and ^37^IAWSTVNKQDGGGKKKSH^54^ (AGH35565.1) **(C-D)** uniquely detect in ethanolamine samples. b/y ions with neutral loss of 59 Da are denoted with *.

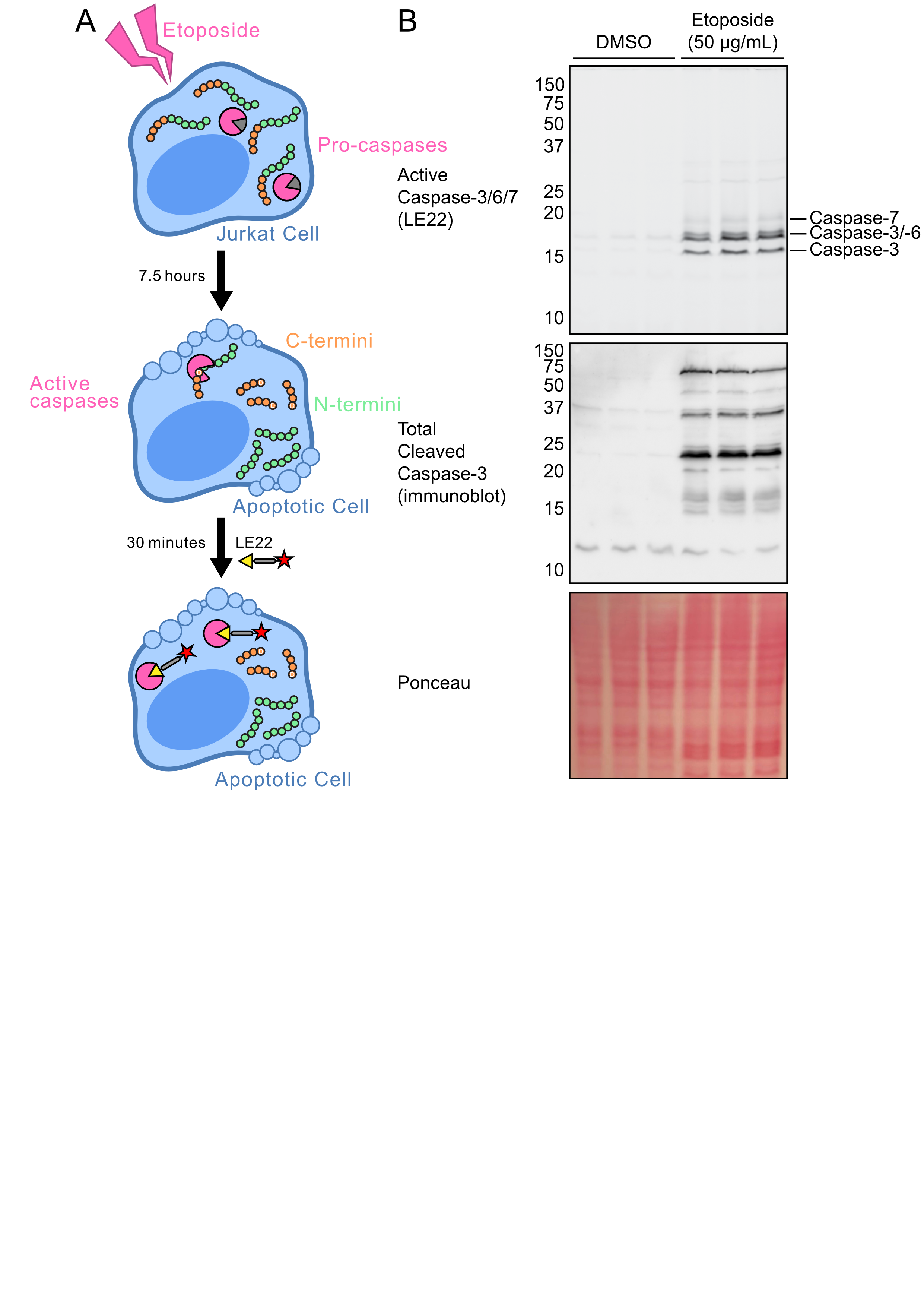
**Figure S6. Induction of apoptosis by etoposide in Jurkat cells. (A)** Model of etoposide-induced apoptosis. Jurkat cells were seeded and treated with 50 µg/mL etoposide for eight hours. Apoptosis induced caspase activation, which cleave peptides following aspartic acid residues. To confirm apoptosis, upregulated caspase-3/-6/-7 activity was measured by live-cell labelling with the activity-based probe LE22 for the final 30 minutes of etoposide treatment. LE22 covalently binds to the active cysteine of caspases for direct analysis of activity by in-gel fluorescence. **(B)** Measurement of active caspases and total cleaved caspase-3 in apoptotic and naïve Jurkat cells. Following lysis and collection, samples were resolved by SDS-PAGE and scanned in-gel for Cy5 fluorescence to detect caspase activity. The gel was transferred for immunoblot analysis of cleaved caspase-3. Ponceau S stain was used as a loading control.

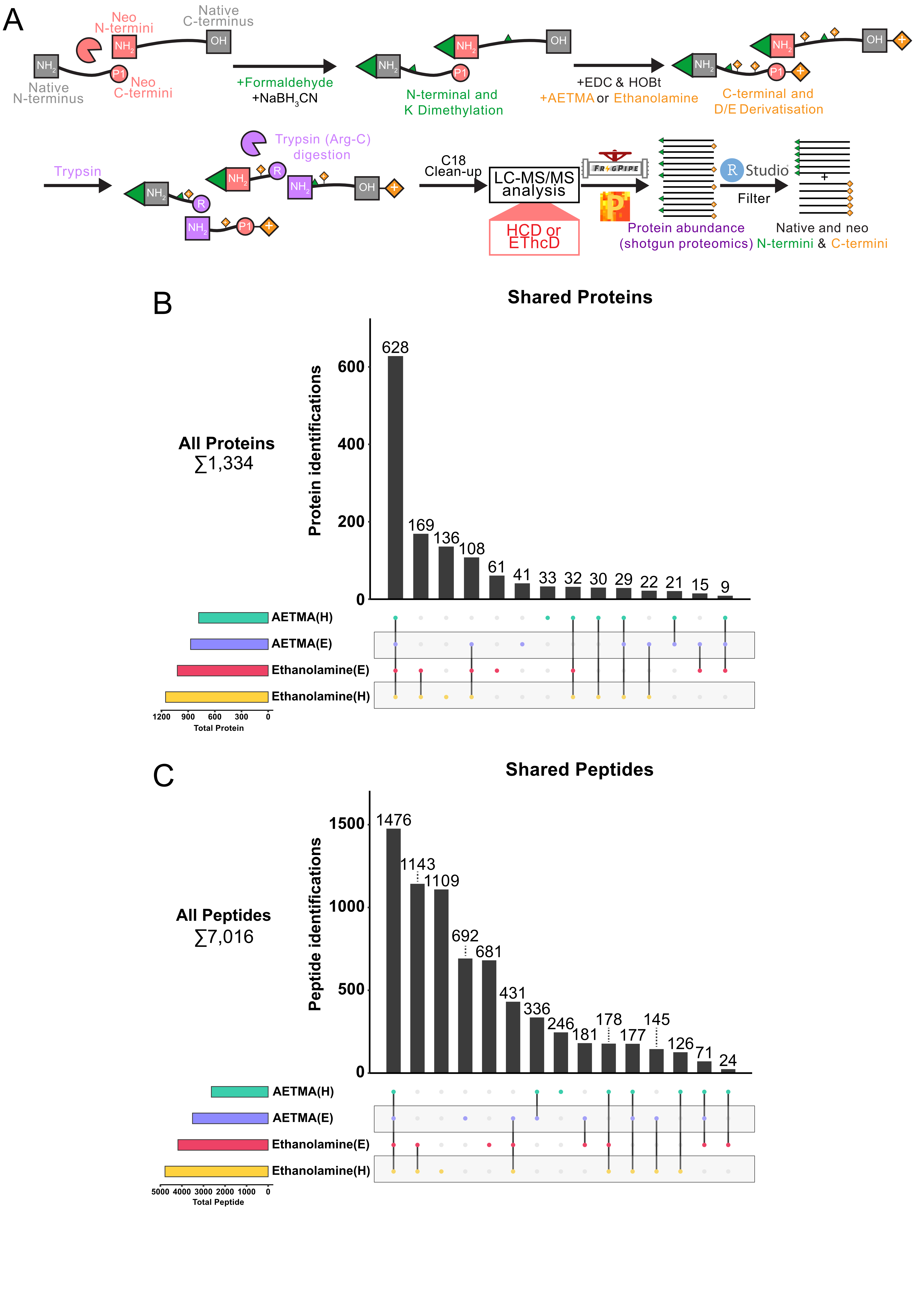

**Figure S7. Overlap of protein and peptide identifications across different labelling and fragmentation strategies.** **(A)** Schematic workflow of carboxyl labelling used for degradomics analysis. Proteolysis results in peptide cleavage, leading to the generation of neo-N-/C-termini (red). Amines were blocked by reductive methylation (green), labelling native and neo-N-termini, as well as lysine residues. Carboxyl groups were then labelled with ethanolamine or AETMA following derivatisation with EDC and HOBt (orange), which labels native and neo-C-termini, as well as aspartate and glutamate residues. Trypsin was used to further digest the sample for LC-MS/MS analysis, generating unlabelled N/C-termini (purple) which were differentiated from native and neo-N/C-termini. Peptides were then subjected to LC-MS/MS analysis with fragmentation by either HCD or EThcD and the resulting data were searched and analysed using FragPipe and Perseus for peptide identification, quantification, and statistics. N/C-termini were filtered based on their respective chemical tags using RStudio. (B-C) Upset plot of overlapping protein **(B)** or total peptide **(C)** identifications between each method.

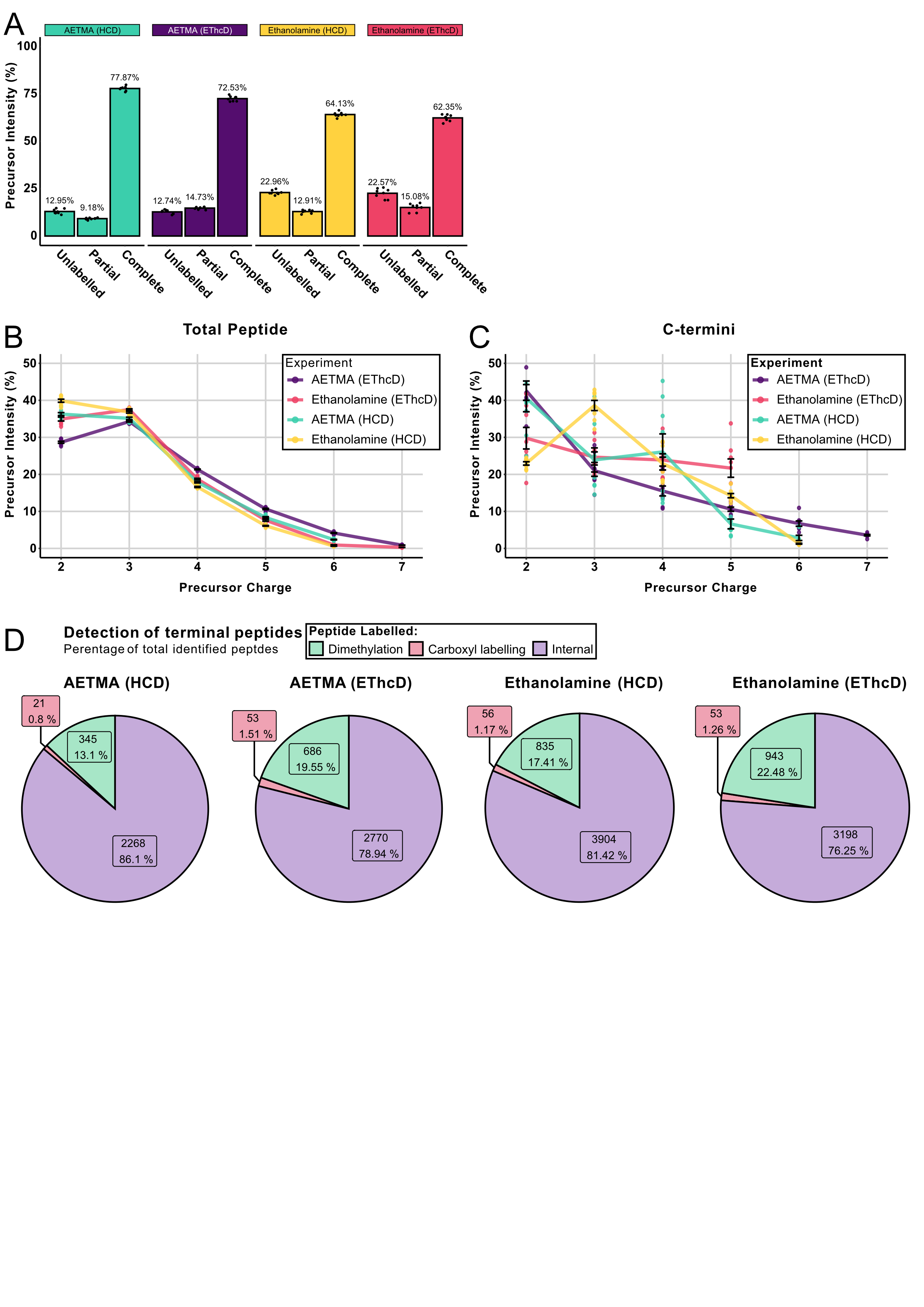

**Figure S8. AETMA increases carboxyl labelling efficiency and precursor charge states.** **(A)** Carboxyl labelling efficiency of identified peptides based on internal aspartate and glutamate residues for each labelling and fragmentation method. Peptides with all aspartate/glutamate residues labelled by AETMA/ethanolamine are complete, whereas those missing labelling on one or more residues are partial or unlabelled, if none are labelled. Graphs are depicted based on the summed percentage intensity of precursors for each group (n = 4/group). **(B-C)** Precursor charges observed for each labelling and fragmentation method at the total peptide **(B)** and C-terminal peptide **(C)** level. Graphs are depicted based on the summed percentage intensity of precursors for each group (n = 4/group). **(D)** Proportion of internal digests (purple), dimethylated N-termini (green), and labelled C-termini (red) within each experiment.

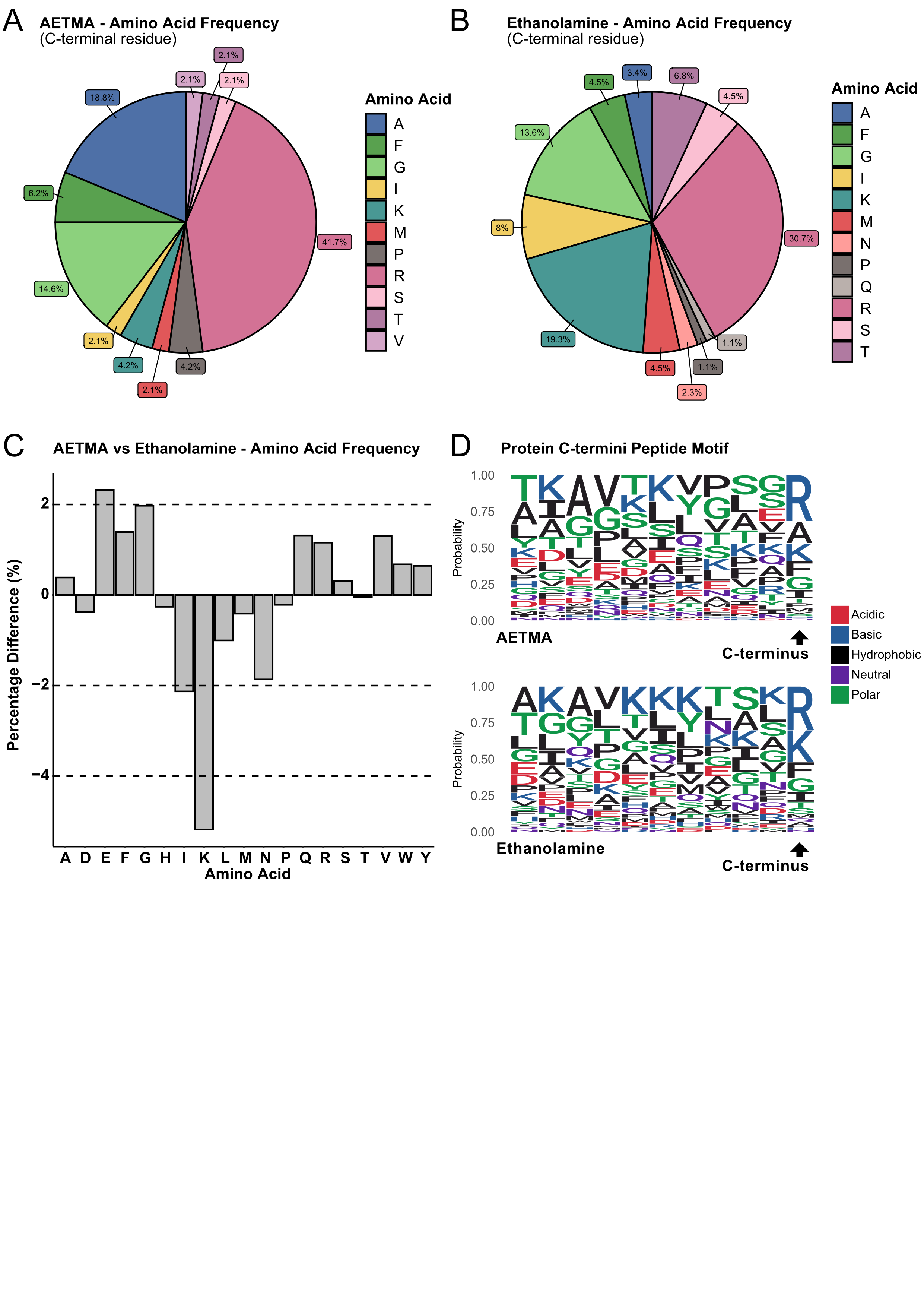

**Figure S9. Amino acid preferences for detected C-termini in Jurkat cells. (A-B)** Frequency of amino acids observed in the total identified C-termini for AETMA **(A)** or ethanolamine **(B)** labelled peptides. **(C)** Amino acid frequency within the identified C-terminal peptides measured as percentage differences between AETMA and ethanolamine labelled samples. Positive values indicate an enrichment of amino acids in AETMA, and depletion for negative values, and vice-versa for ethanolamine. **(D)** Peptide sequence logos of unique C-termini identified following labelling with either AETMA or ethanolamine. The final position is considered the identified C-terminus.
